# Neural Correlates of Psychedelic, Sleep, and Sedated States Support Global Theories of Consciousness

**DOI:** 10.1101/2024.10.23.619731

**Authors:** Rui Dai, Hyunwoo Jang, Anthony G. Hudetz, Zirui Huang, George A. Mashour

## Abstract

Understanding neural mechanisms of consciousness remains a challenging question in neuroscience. A central debate in the field concerns whether consciousness arises from global interactions that involve multiple brain regions or focal neural activity, such as in sensory cortex. Additionally, global theories diverge between the Global Neuronal Workspace (GNW) hypothesis, which emphasizes frontal and parietal areas, and the Integrated Information Theory (IIT), which focuses on information integration within posterior cortical regions. To disentangle the global vs. local and frontoparietal vs. posterior dilemmas, we measured global functional connectivity and local neural synchrony with functional magnetic resonance imaging (fMRI) data across a spectrum of conscious states in humans induced by psychedelics, sleep, and deep sedation. We found that psychedelic states are associated with increased global functional connectivity and decreased local neural synchrony. In contrast, non-REM sleep and deep sedation displayed the opposite pattern, suggesting that consciousness arises from global brain network interactions rather than localized activity. This mirror-image pattern between enhanced and diminished states was observed in both anterior-posterior (A-P) and posterior-posterior (P-P) brain regions but not within the anterior part of the brain alone. Moreover, anterior transmodal regions played a key role in A-P connectivity, while both posterior transmodal and posterior unimodal regions were critical for P-P connectivity. Overall, these findings provide empirical evidence supporting global theories of consciousness in relation to varying states of consciousness. They also bridge the gap between two prominent theories, GNW and IIT, by demonstrating how different theories can converge on shared neuronal mechanisms.

## Introduction

Consciousness has long been regarded as a philosophical “hard problem” due to the challenge of explaining how subjective experience arises from the brain (Chalmers, 1995). Since the 1990s, advances in neuroimaging technologies—particularly functional magnetic resonance imaging (fMRI)—have enabled the observation of brain activity and its linkage to various cognitive and conscious states, facilitating the search for the neural correlates of consciousness (Crick and Koch, 1990; Koch et al., 2016). These developments have transformed the neuroscientific exploration of consciousness into a vibrant field, encompassing a wealth of empirical research and inspiring numerous theoretical frameworks (Crick and Koch, 2003; Dehaene and Changeux, 2011; Koch et al., 2016; Mashour and Hudetz, 2018; Seth and Bayne, 2022; Storm et al., 2024; Yaron et al., 2022). One debate in this field is whether consciousness arises from a global process involving widespread brain networks or from localized neural activity, e.g., in sensory cortices. Prominent global theories, such as the Global Neuronal Workspace (GNW) hypothesis (Dehaene and Changeux, 2011; Dehaene and Naccache, 2001; Mashour et al., 2020) and Integrated Information Theory (IIT) (Tononi, 2008, 2004; Tononi et al., 2016) argue that consciousness emerges from networks that extend beyond sensory cortex but they differ markedly in terms of the role of the prefrontal cortex. GNW highlights the global broadcast of information, particularly between the frontal and parietal regions (Dehaene and Changeux, 2004; Dehaene and Naccache, 2001). On the other hand, IIT suggests that consciousness emerges from complex interactions, likely residing in a posterior cortical “hot zone” (Koch et al., 2016; Oizumi et al., 2014; Tononi, 2004). We refer to both GNW and IIT as ‘global’ theories because consciousness is proposed to be a function of complex interactions across large-scale brain networks rather than generated within a single defined brain region. In contrast, local theories propose that sensory cortex activity is sufficient for conscious experience. For instance, the Recurrent Processing Theory (RPT) posits that horizontal connections and recurrent processing within sensory areas are critical for consciousness (Lamme et al., 2001; Lamme, 2010, 2006).

In this context of competing theories, a critical question arises: which theoretical framework offers the most accurate explanation of consciousness? One approach is to employ adversarial testing, where theories are evaluated based on differential, empirically testable predictions (Melloni et al., 2023). However, most existing research has focused on testing individual theories independently, with few studies directly comparing multiple competing frameworks. Additionally, traditional studies have primarily concentrated on isolated states, such as sleep, anesthesia, or coma, often employing distinct theoretical frameworks and methodologies. To address this gap, our study adopts an integrative approach to assessing select theories of consciousness by examining a broad spectrum of conscious states, encompassing physiological and pharmacological conditions, from psychedelic experiences to natural sleep and varying levels of sedation. We implemented a comprehensive set of methodologies that focus on empirically driven core features represented in both global and local theories of consciousness. This includes well-established techniques such as functional connectivity and regional homogeneity, as well as advanced methods like the topological integration (measured by global efficiency) and machine learning-based feature importance analysis. Specifically, by analyzing fMRI data across psychedelic states of consciousness (induced by LSD, as well as subanesthetic ketamine and nitrous oxide) and diminished states of consciousness (induced by sleep or deep sedation with propofol), we aim to address two fundamental questions: (1) Does a global or local framework for consciousness provide a more compelling explanation for the observed variations in the neural correlates of conscious states? and (2) If global theories are supported, is GNW or IIT a more accurate representation?

In this study, we employed between-network functional connectivity as a global metric, and regional homogeneity as a local metric. If consciousness is primarily a global phenomenon, we would expect to see increased connectivity across brain networks during psychedelic states and a breakdown of global connectivity during states of diminished consciousness like sleep and deep sedation. Conversely, if consciousness is primarily local, we would anticipate heightened local neural synchrony during psychedelic states and local decoherence during sleep or deep sedation (Figure 1a). To further compare and evaluate the two network-based approaches, GNW and IIT, we not only conducted conventional functional connectivity analysis but also applied a graph-theoretical metric (specifically, global efficiency) to quantify functional integration (Jang et al., 2024b). We analyzed the brain’s functional connectivity along both its anatomical axis (from anterior to posterior regions) and its functional axis (from unimodal sensory to transmodal integrative regions). If the GNW framework is accurate, we would predict increased functional connectivity between anterior and posterior regions during psychedelic states, and a decrease in this connectivity during sleep or sedative states. On the other hand, if IIT is more accurate, we would predict a primary increase in functional connectivity within posterior regions during psychedelic states, and a decrease in connectivity within these same regions during dreamless sleep or anesthesia (Figure 1b).

**Figure 1.**
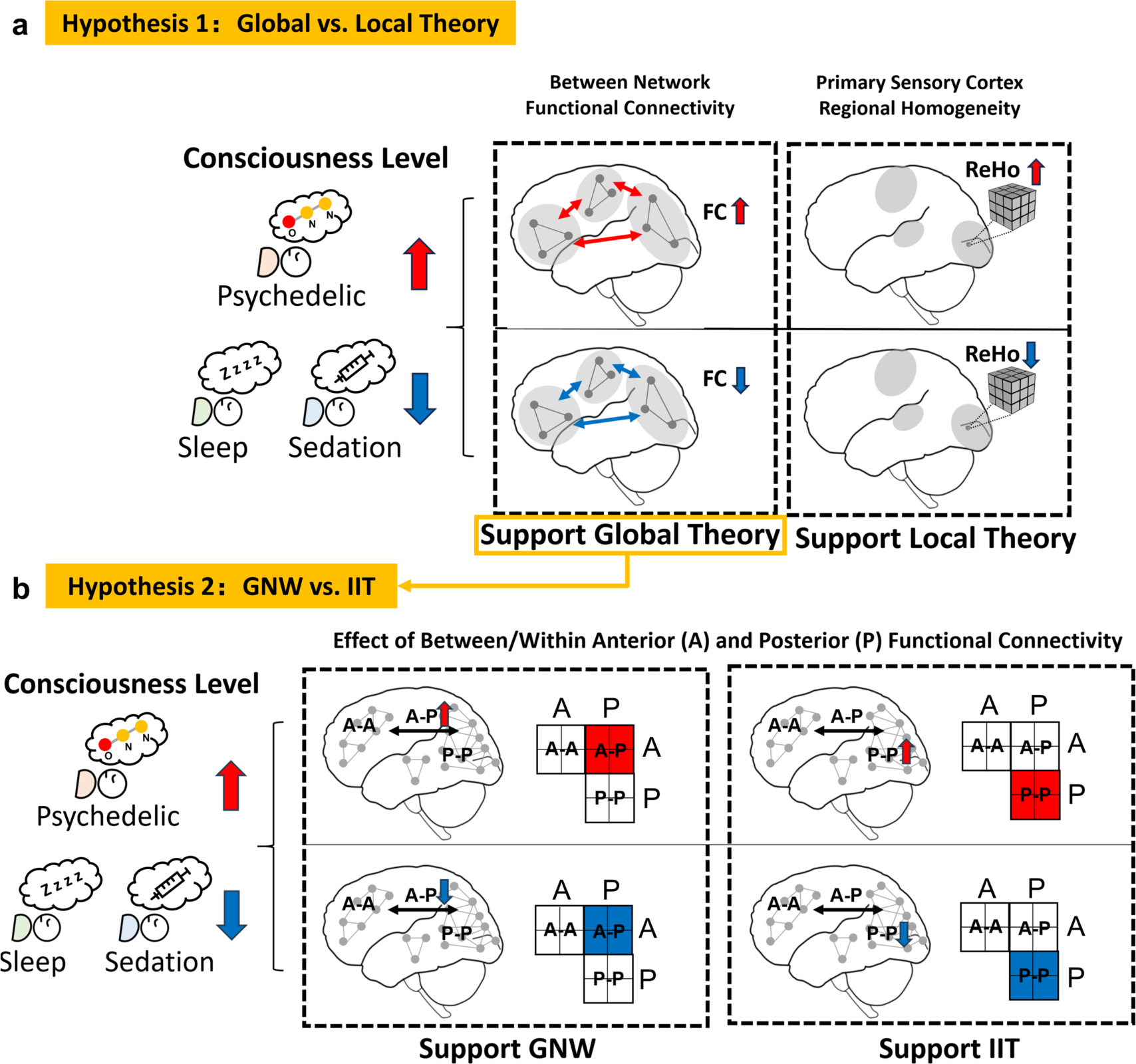
Theoretical Predictions of Global and Local Connectivity in Different States of Consciousness. **(a)** The expected outcomes for global and local theories of consciousness. The global theory predicts increased between-network functional connectivity (FC) in heightened consciousness states (e.g., psychedelics) and decreased FC in diminished states (e.g., sleep or sedation). The local theory predicts the opposite pattern for local connectivity (ReHo). **(b)** The expected outcomes for Global Neuronal Workspace (GNW) and Integrated Information Theory (IIT) related to anterior-posterior (A-P) and posterior-posterior (P-P) functional connectivity across different states of consciousness. Increased A-P connectivity during psychedelic states and decreased A-P connectivity during sleep and sedative states would support GNW. Conversely, increased P-P connectivity during psychedelic states and decreased P-P connectivity during sleep and sedative states would support IIT.

## Results

We investigated three classical and non-classical psychedelic agents—LSD (n=15), ketamine (n=12), and nitrous oxide (n=15)—all known to induce non-ordinary states of consciousness that can be considered phenomenologically enriched. Sleep and sedation, which are associated with diminished consciousness, served as comparative conditions. For sleep, we analyzed non-REM sleep stages N1; n=33 and N2; n=29. For sedation, we examined the effects of propofol at various effect-site concentrations: 1.0 μg/ml (n=12), 1.9 μg/ml (n=12), 2.4 μg/ml (n=25), and 2.7 μg/ml (n=26). Each altered state of consciousness was meticulously compared to its corresponding baseline condition to assess changes in functional connectivity.

We initiated our analysis by examining between-network functional connectivity, a global measure of brain activity that gauges the correlation between distinct brain networks, including default-mode, frontoparietal, limbic, ventral attention, dorsal attention, somatosensory, and visual networks. Our results demonstrated an overall increase in between-network functional connectivity across all psychedelic conditions (Effect sizes were calculated using Cohen’s d; LSD: d = 0.65, pFDR = 0.043; ketamine: d = 0.78, pFDR = 0.041; nitrous oxide: d = 0.60, pFDR = 0.049), suggesting heightened global brain integration during these altered states. In contrast, during non-REM sleep stage 2, we observed a significant decrease in between-network functional connectivity compared to baseline (d = -0.56, pFDR = 0.017), suggesting a diminished state of global integration. Similarly, under varying doses of propofol, a consistent decrease in between-network functional connectivity was evident (propofol 1.9 μg/ml: d = -0.73, pFDR = 0.045; 2.4 μg/ml: d = -0.77, pFDR = 0.003; 2.7 μg/ml: d = -1.49, pFDR = 0.0006), further supporting the hypothesis of reduced global brain integration in states of diminished consciousness (Figure 2a and 2b; Figure S1). Effect sizes were calculated using Cohen’s d.

**Figure 2.**
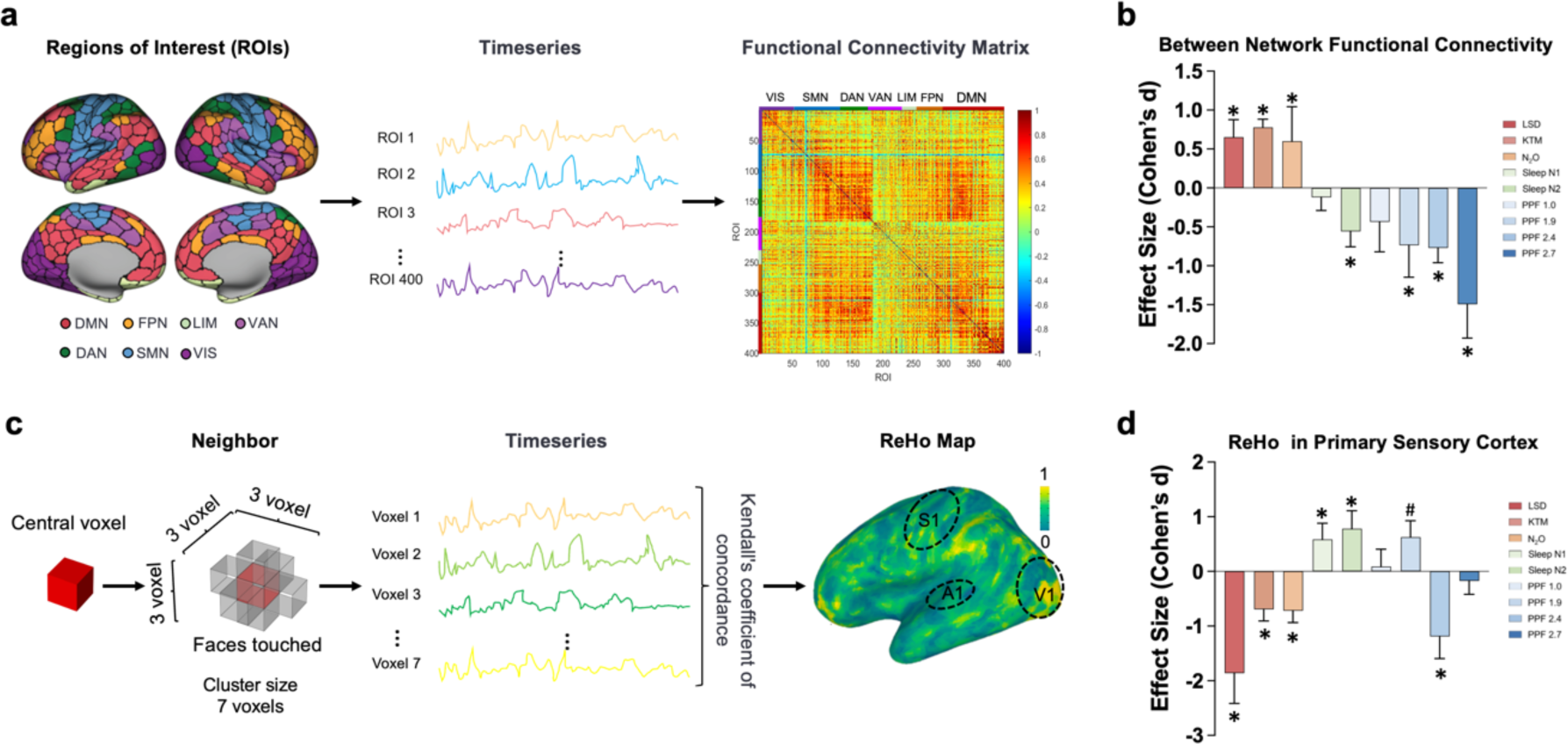
Global and Local Connectivity in Different States of Consciousness. **(a)** Schematic of between-network functional connectivity. A total of 400 regions of interest (ROIs) were defined across the brain, spanning seven established functional networks: default-mode (DMN), frontoparietal (FPN), limbic (LIM), ventral attention (VAN), dorsal attention (DAN), somatomotor (SMN), and visual (VIS). Time series data from each ROI were used to calculate between-network functional connectivity by correlating ROIs in one network with those in the other six. The resulting values were averaged to produce a single between-network functional connectivity for each network, and these network-specific values were further averaged to derive a global between-network functional connectivity for the entire brain. **(b)** Effect size of between-network functional connectivity. The bar graph displays the effect sizes for between-network functional connectivity across various conscious states. Statistical significance was determined using paired-sample t-tests based on Fisher z-transformed functional connectivity values. **(c)** Schematic of regional homogeneity (ReHo) analysis. This analysis quantifies the similarity of time series data among neighboring brain voxels. ReHo values, representing the level of synchronization within a given brain area, are extracted from primary sensory cortices: the primary somatosensory cortex (S1), primary auditory cortex (A1), and primary visual cortex (V1). **(d)** Effect Size of ReHo in the primary sensory cortices. This bar graph shows the effect sizes for ReHo in the primary sensory cortex across different states of consciousness. Statistical significance was assessed using paired-sample t-tests on Fisher z-transformed ReHo values. Significant differences from baseline are marked by asterisks (*, FDR-corrected p < 0.05), and significant differences before FDR correction are indicated by a hash (#, uncorrected p < 0.05). LSD: lysergic acid diethylamide, KTM: ketamine, N2O: nitrous oxide, SLEEP N1: non-REM sleep stage 1, SLEEP N2: non-REM sleep stage 2, PPF 1.0: propofol 1.0 μg/ml, PPF 1.9: propofol 1.9 μg/ml, PPF 2.4: propofol 2.4 μg/ml, PPF 2.7: propofol 2.7 μg/ml.

Next, we conducted a regional homogeneity (ReHo) (Zang et al., 2004) analysis to investigate changes in local connectivity. ReHo analysis quantifies the synchronization of neural activity within localized brain regions by calculating Kendall’s W for each voxel within a 7-voxel neighborhood. The resulting voxel-level W values are then aggregated to produce region-specific ReHo measures, such as those for primary sensory cortices. Our analysis of the psychedelic datasets consistently showed a decrease in ReHo (LSD: d = - 1.86, pFDR < 0.0001; ketamine: d = -0.69, pFDR = 0.049; nitrous oxide: d = -0.72, pFDR = 0.033), indicating reduced local synchrony of neural activity. In contrast, the datasets for sleep and propofol generally exhibited an increase in ReHo (Sleep N1: d = 0.59, pFDR = 0.006; Sleep N2: d = 0.78, pFDR = 0.0009; propofol 1.9 μg/ml: d = -0.63, uncorrected p = 0.049), suggesting enhanced local synchrony (except for the high doses of propofol) (Figure 2c and 2d, Figure S2). These results do not support local theory, which predicts increased local connectivity during psychedelic states and decreased synchrony during states of diminished consciousness. Instead, the observed patterns suggest that local connectivity alone may not fully explain the changes in conscious states.

With between-network functional connectivity and ReHo data supporting global approaches to consciousness, we focused our attention on global theories and assessed whether GNW or IIT provides a more compelling explanation for the changes in conscious state. While GNW highlights the importance of frontal and parietal connections, and IIT emphasizes posterior cortical areas, neither theory offers a precise delineation of the specific brain regions implicated in their frameworks. Given this lack of clarity, we strategically employed an anatomical division of the brain into two parts based on the central sulcus: the anterior region, which includes the frontal lobe, and the posterior region, encompassing the parietal, temporal, and occipital lobes. We then assessed functional connectivity between anterior and posterior (A-P) brain regions, as well as within anterior (A-A) and within posterior (P-P) regions (Figure 3a). Effect sizes were calculated using Cohen’s d. In psychedelic states, we observed a significant increase in P-P functional connectivity (LSD: d = 0.62, pFDR= 0.041; ketamine: d = 0.76, pFDR = 0.038; nitrous oxide: d = 0.74, pFDR = 0.028), followed by enhanced A-P functional connectivity for non-classical psychedelics (ketamine: d = 0.87, pFDR = 0.026; nitrous oxide: d = 0.60, uncorrected p = 0.037).

**Figure 3.**
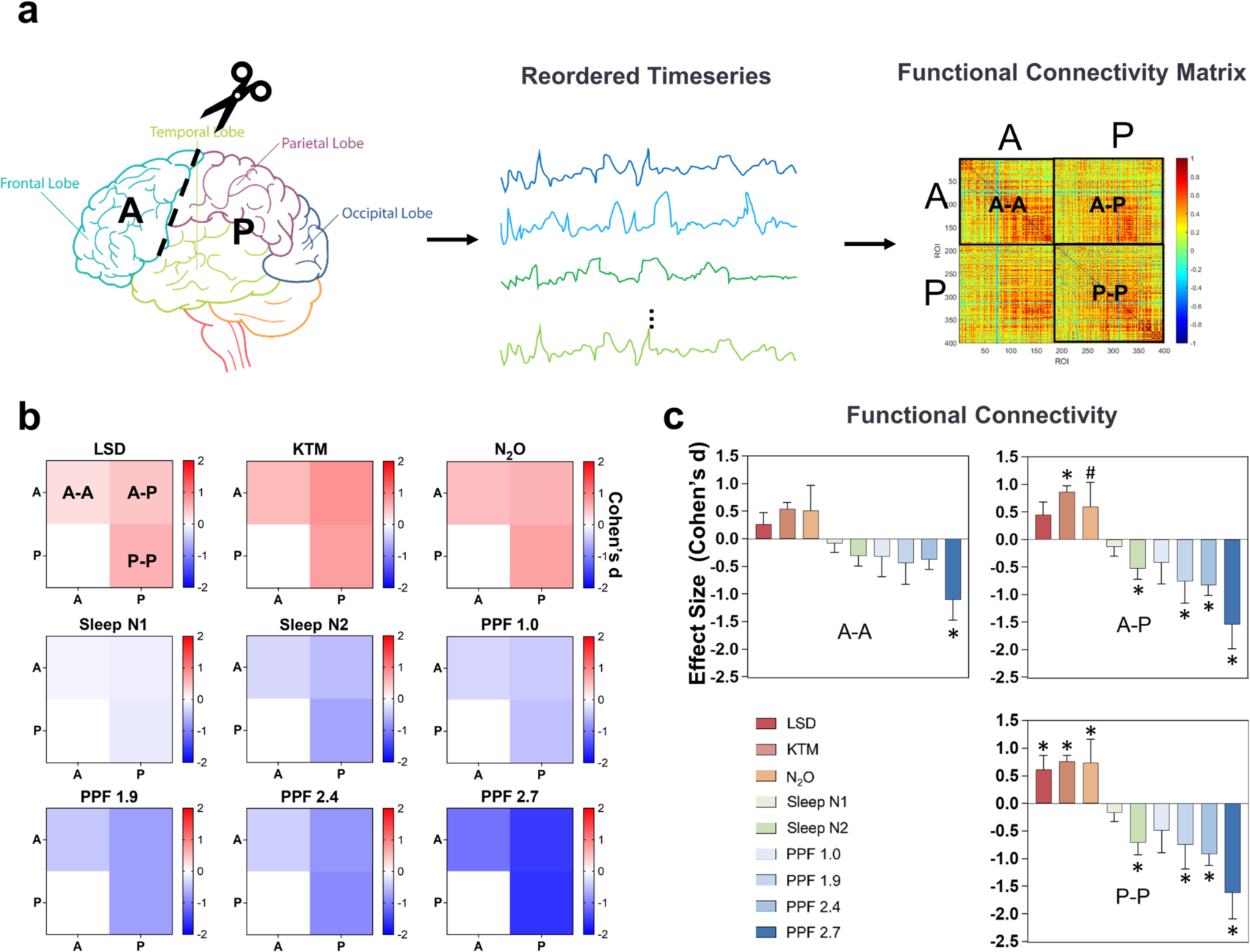
Anterior vs. Posterior Connectivity in Different States of Consciousness. **(a)** Schematic of the analysis pipeline. Functional connectivity was assessed by dividing the brain into anterior (A) and posterior (P) regions based on the central sulcus, with time series extracted from regions of interest (ROIs) across the brain. Functional connectivity matrices were constructed by calculating pairwise connectivity between all ROIs within anterior regions (A-A), within posterior regions (P-P), and between anterior and posterior regions (A-P), followed by averaging the connectivity values within each category. **(b)** Heat maps display the effect size (Cohen’s d) of functional connectivity changes between each altered state of consciousness and its corresponding baseline for A-A, A-P, and P-P regions. Warmer colors (red) represent increased connectivity, while cooler colors (blue) indicate reduced connectivity. Each entry in the 2×2 matrix reflects the effect size of functional connectivity between the specified region pairs in the given state versus its baseline. **(c)** Effect size (Cohen’s d) of functional connectivity changes in different states of consciousness. Bar plots display the effect sizes of functional connectivity changes under different conditions, including LSD, ketamine (KTM), nitrous oxide (N2O), sleep (N1 and N2), and propofol (PPF) at various infusion rates, compared to their respective baselines across A-A, A-P, and P-P regions. Statistical significance was determined using paired-sample t-tests based on Fisher z-transformed functional connectivity values. Significant differences from baseline are marked by asterisks (*, FDR-corrected p < 0.05), and significant differences before FDR correction are indicated by a hash (#, uncorrected p < 0.05).

In contrast, during sleep and deep sedation, we found pronounced decreases in both A-P functional connectivity (sleep N2: d = -0.53, pFDR= 0.024; propofol 1.9 μg/ml: d = -0.76, pFDR= 0.041; propofol 2.4 μg/ml: d = -0.83, pFDR = 0.002; propofol 2.7 μg/ml: d = -1.55, pFDR < 0.0001) and P-P functional connectivity (sleep N2: d = -0.71, pFDR = 0.002; propofol 1.9 μg/ml: d = -0.75, pFDR= 0.038; propofol 2.4 μg/ml: d = -0.91, pFDR = 0.0005; propofol 2.7 μg/ml: d = -1.16, pFDR < 0.0001) (Figure 3b, c, and Figure S3). Notably, there was no statistical significance for changes in A-A functional connectivity, suggesting that functional connectivity in frontal cortex alone does not fully account for variations in conscious state.

To achieve a more refined understanding beyond the anatomical axis from anterior to posterior cortical regions, we further subdivided these brain regions based on a functional axis ranging from unimodal (sensory and motor cortices) to transmodal (association cortices) processing (Margulies et al., 2016). Regions with gradient values greater than 0 were classified as transmodal, and those with values less than 0 as unimodal (see Methods for details). We then analyzed functional connectivity between and within anterior transmodal (AT), anterior unimodal (AU), posterior transmodal (PT), and posterior unimodal (PU) regions (Figure 4a). Overall, opposite changes were observed across the two classes of states, with increased functional connectivity during psychedelic states and decreased connectivity during sleep and deep sedation (Figure 4b, c, and Figure S4). A consistent ‘mirror-image’ pattern in functional connectivity was observed, particularly in the AT-PU and PT-PU regions, where connectivity significantly increased during psychedelic states and decreased during sedative-hypnotic states compared to each baseline. The effect sizes, measured using Cohen’s d, were as follows: For AT-PU, LSD: d = 0.95, pFDR = 0.009; ketamine: d = 0.92, pFDR = 0.019; nitrous oxide: d = 0.76, pFDR = 0.019; sleep N2: d = -0.43, pFDR = 0.036; propofol 1.9 μg/ml: d = -0.75, pFDR = 0.036; propofol 2.4 μg/ml: d = -0.60, pFDR = 0.019; propofol 2.7 μg/ml: d = -1.31, pFDR < 0.0001. For PT-PU, LSD: d = 1.01, pFDR = 0.006; ketamine: d = 0.84, pFDR = 0.021; nitrous oxide: d = 0.91, pFDR = 0.008; sleep N2: d = -0.64, pFDR = 0.006; propofol 2.4 μg/ml: d = -0.61, pFDR = 0.011; propofol 2.7 μg/ml: d = -1.41, pFDR < 0.0001. A-P regions, such as AT-PT and AU-PT, along with P-P regions like PT-PT and PU-PU, displayed similar trends of connectivity, but these lacked consistent statistically significant differences.

**Figure 4.**
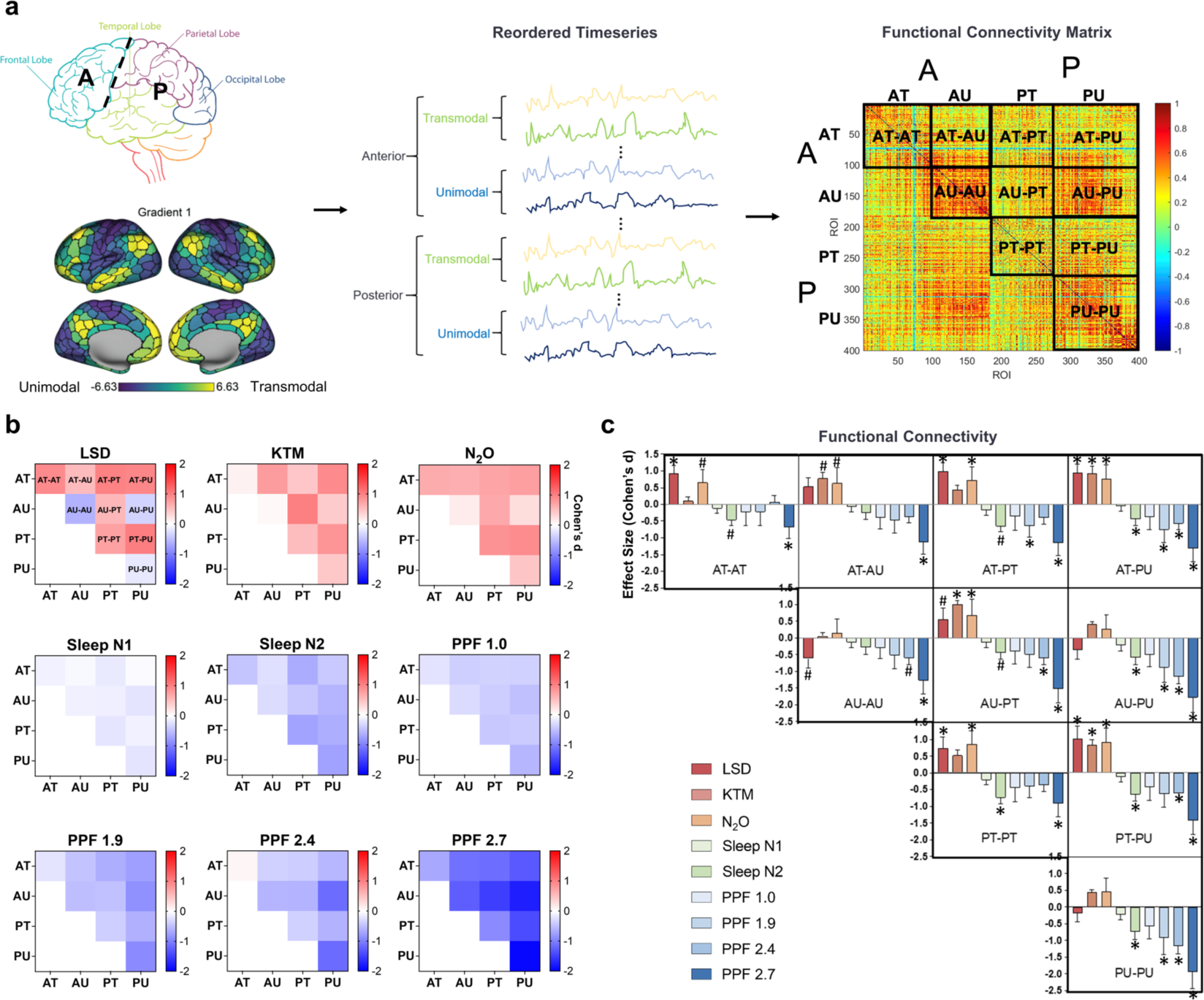
Anterior/Posterior vs. Transmodal/Unimodal Functional Connectivity: **(a)** Analytical pipeline for connectivity assessment. The brain was first divided into anterior (A) and posterior (P) regions based on anatomical structure using the central sulcus as the boundary. These anterior and posterior regions were then further subdivided based on functional gradients, separating transmodal (higher-order cognitive) areas from unimodal (sensory-motor) areas. Time series were extracted from regions of interest (ROIs) across these divisions, and functional connectivity matrices were calculated between and within anterior transmodal (AT), anterior unimodal (AU), posterior transmodal (PT), and posterior unimodal (PU) regions. **(b)** Heatmaps show the effect size of functional connectivity changes during different states. Each entry in the 4×4 heatmap represents the effect size of functional connectivity between pairs of brain regions: AT, AU, PT, and PU. Red hues indicate increased connectivity, while blue hues indicate reduced connectivity. **(c)** Bar plots display effect sizes of functional connectivity changes under different conditions: LSD, ketamine (KTM), nitrous oxide (N2O), sleep (N1 and N2), and propofol at various effect-site concentrations. Statistical significance was determined using paired-sample t-tests based on Fisher z-transformed FC values. Significant differences from baseline are marked by asterisks (*, FDR-corrected p < 0.05), and significant differences before FDR correction are indicated by a hash (#, uncorrected p < 0.05).

To provide a comprehensive depiction of anterior vs. posterior and unimodal vs. transmodal functional connectivity, we created a circular plot highlighting all connections with strong effect sizes (|Cohen’s d| > 0.8) across all datasets. In this visualization, regions are arranged from anterior transmodal to posterior unimodal areas, with each region labeled according to its functional network and anatomical location in the brain. Additionally, we conducted a meta-analytic approach of cognitive functions associated with each of the AT, AU, PT, and PU regions to provide insights into the functional roles of brain region groups. We then examined connectivity density by counting the number of connections (i.e., edges) between or within region groups. During psychedelic states, we observed a notable increase in connectivity density, particularly for A-P functional connectivity and P-P functional connectivity, irrespective of the unimodal vs. transmodal specialization of the areas involved. In contrast, during sleep and sedation, a marked decrease in connectivity density was most pronounced in A-P and P-P regions. Additionally, we conducted a detailed analysis of edge ratios, which quantify the proportion of robust connectivity changes relative to the maximum possible connections between and within anterior and posterior regions, including both transmodal and unimodal areas. Our findings indicate that during psychedelic states, edge ratios were elevated in A-P and P-P regions, suggesting increased connectivity. Conversely, during deep sleep and sedation, edge ratios were reduced, particularly in the A-P and P-P regions (Figure 5), underscoring the mirror-image pattern between enhanced and diminished states of consciousness.

**Figure 5.**
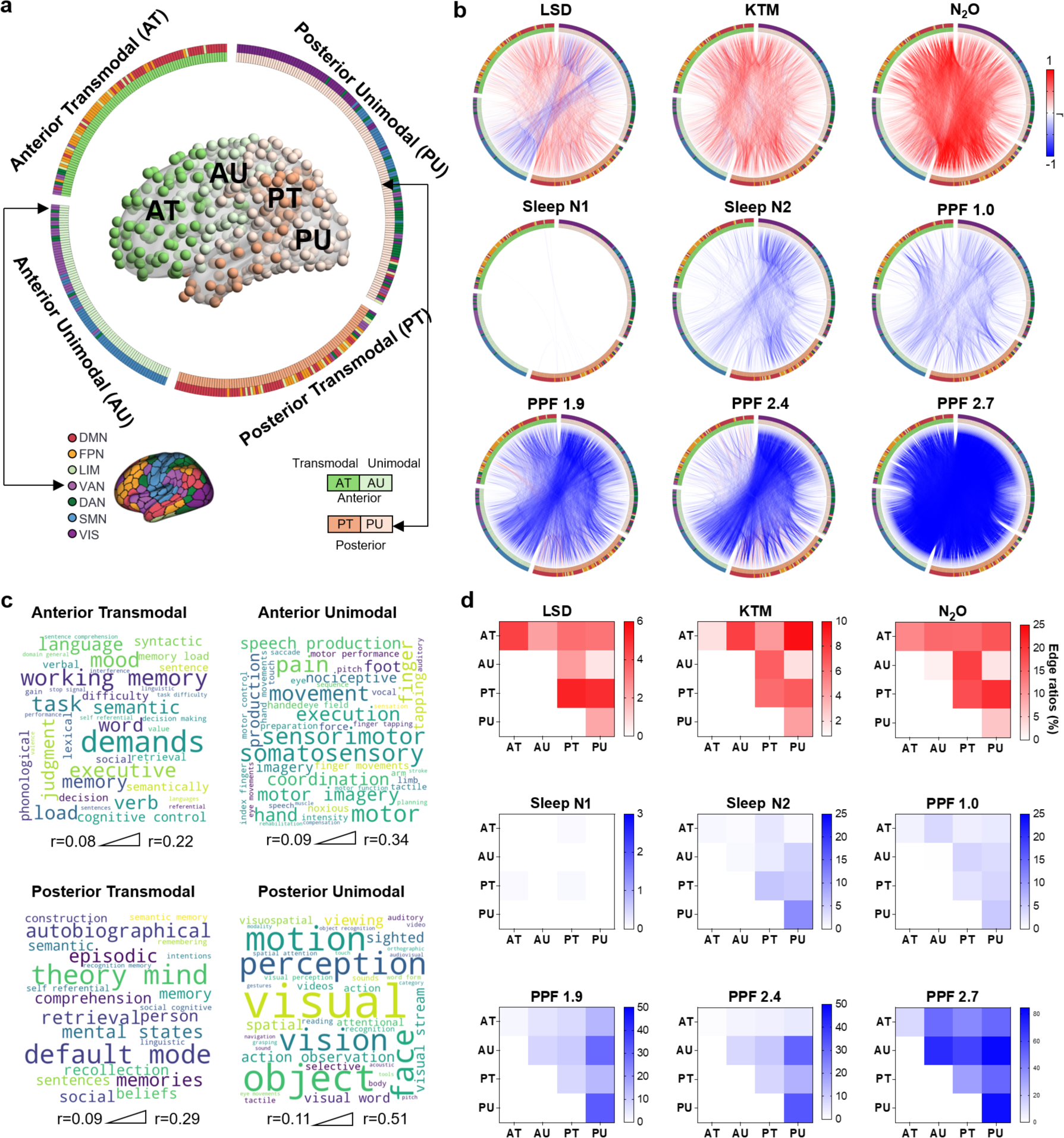
Functional Connectivity Density and Cognitive Correlates Across Anterior-Posterior and Transmodal-Unimodal Brain Regions in Different States of Consciousness. **(a)** The brain map illustrates the spatial distribution of regions categorized as anterior transmodal (AT), anterior unimodal (AU), posterior transmodal (PT), and posterior unimodal (PU). The inner ring of the circular plot indicates whether each region is unimodal or transmodal, while the outer ring specifies the functional network to which the regions belong. **(b)** Circular Plots of Connectivity Changes: Circular plots present the connectivity changes under various conditions, compared to their respective baselines. Each connection (i.e., edge) represents a strong effect size (|effect size| > 0.8). **(c)** Word Clouds of Cognitive Functions: Word clouds depict cognitive functions associated with brain regions, with font sizes proportional to the correlation coefficient (r) between the anatomical mask and the meta-analytic map generated by Neurosynth (https://www.neurosynth.org/) for a given term. The triangle beneath the word clouds indicates the relationship between font size and correlation coefficient. **(d)** Edge ratio (% of maximum possible connections): Heatmaps illustrate the quantified edge ratios, defined as the ratio of robust connectivity changes to the maximum possible connections. LSD: lysergic acid diethylamide, KTM: ketamine, N2O: nitrous oxide, Sleep N1: non-REM sleep stage 1, Sleep N2: non-REM sleep stage 2, PPF 1.0: propofol 1.0 μg/ml, PPF 1.9: propofol 1.9 μg/ml, PPF 2.4: propofol 2.4 μg/ml, PPF 2.7: propofol 2.7 μg/ml. DMN: default-mode network, FPN: frontoparietal network, LIM: limbic network, VAN: ventral attention network, DAN: dorsal attention network, SMN: somatomotor network, VIS: visual network.

In addition to the classical functional connectivity method, we incorporated an approach from graph theory to further assess the topological integration between and within anterior and posterior regions (Jang et al., 2024b). Specifically, we calculated the normalized multi-level global efficiency (see Methods for detailed definitions), indexing the degree of relative topological integration, between and within anterior and posterior regions (Figure 6, Figure S5, and S6). Consistent with the functional connectivity results, we observed increased A-P and P-P efficiency during psychedelic states and decreased A-P and P-P efficiency during sleep and deep sedation. Within these areas, the AU-PU efficiency, as well as all within-posterior efficiency (PT-PU, PT-PT, PU-PU), exhibited the most pronounced changes, suggesting differential engagement of these regions across states of consciousness.

**Figure 6.**
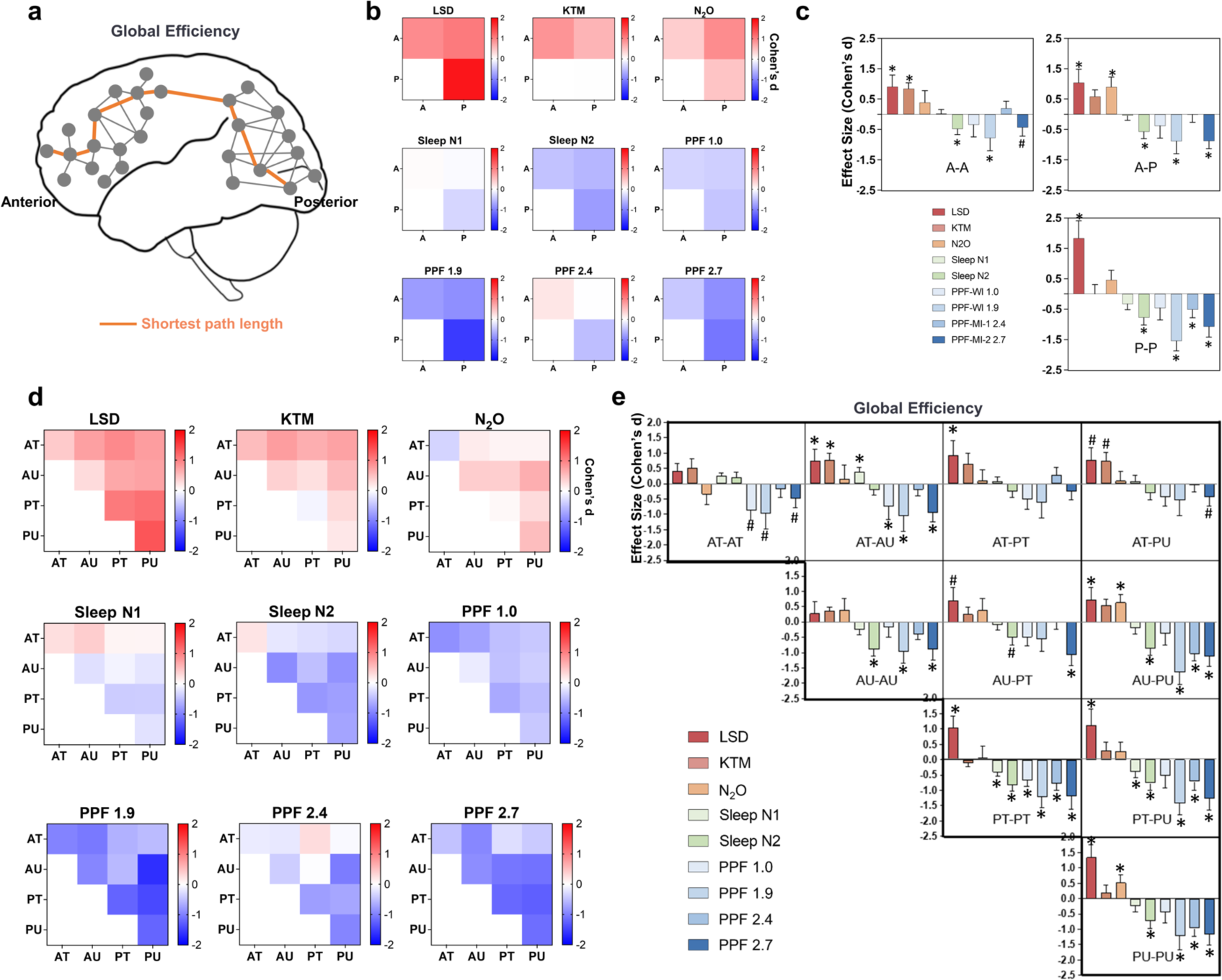
Topological Analysis of Anterior and Posterior Brain Integration Across Different States of Consciousness. **(a)** Schematic of Global Efficiency (Inverse of Shortest Path Length): This illustration demonstrates the calculation of the global efficiency, represented by the average of the inverse of shortest path length between two brain regions. This approach is derived from network science and is a surrogate for the efficiency of information transfer. As an example, orange lines indicate a shortest path between a given pair of brain regions. **(b)** Heatmaps of Global Efficiency: These heatmaps visualize changes in the global efficiency within anterior regions (A-A), between anterior and posterior regions (A-P), and within posterior regions (P-P) regions. Each entry in the heatmap represents the change in global efficiency value between brain regions for a given state relative to its baseline. **(c)** Effect Sizes of Global Efficiency: Bar plots present the effect sizes for changes in the efficiency in A-A, A-P, and P-P regions across various conditions (LSD, KTM, N2O, Sleep N1, Sleep N2, Propofol 1.0, Propofol 1.9, Propofol 2.4, Propofol 2.7). **(d)** Heatmaps of global efficiency between and within anterior transmodal (AT), anterior unimodal (AU), posterior transmodal (PT), and posterior unimodal (PU) regions during different states of consciousness. Each entry in the heatmap represents the change in global efficiency value between brain regions for a given state relative to its baseline. **(e)** Effect sizes for global efficiency across AT, AU, PT, PU regions. Statistical analyses were performed for global efficiency values, with Cohen’s d effect sizes reported to illustrate the magnitude of changes. Significant differences from baseline are marked by asterisks (*, FDR-corrected p < 0.05), and significant differences before FDR correction are indicated by a hash (#, uncorrected p < 0.05).

**Figure 7.**
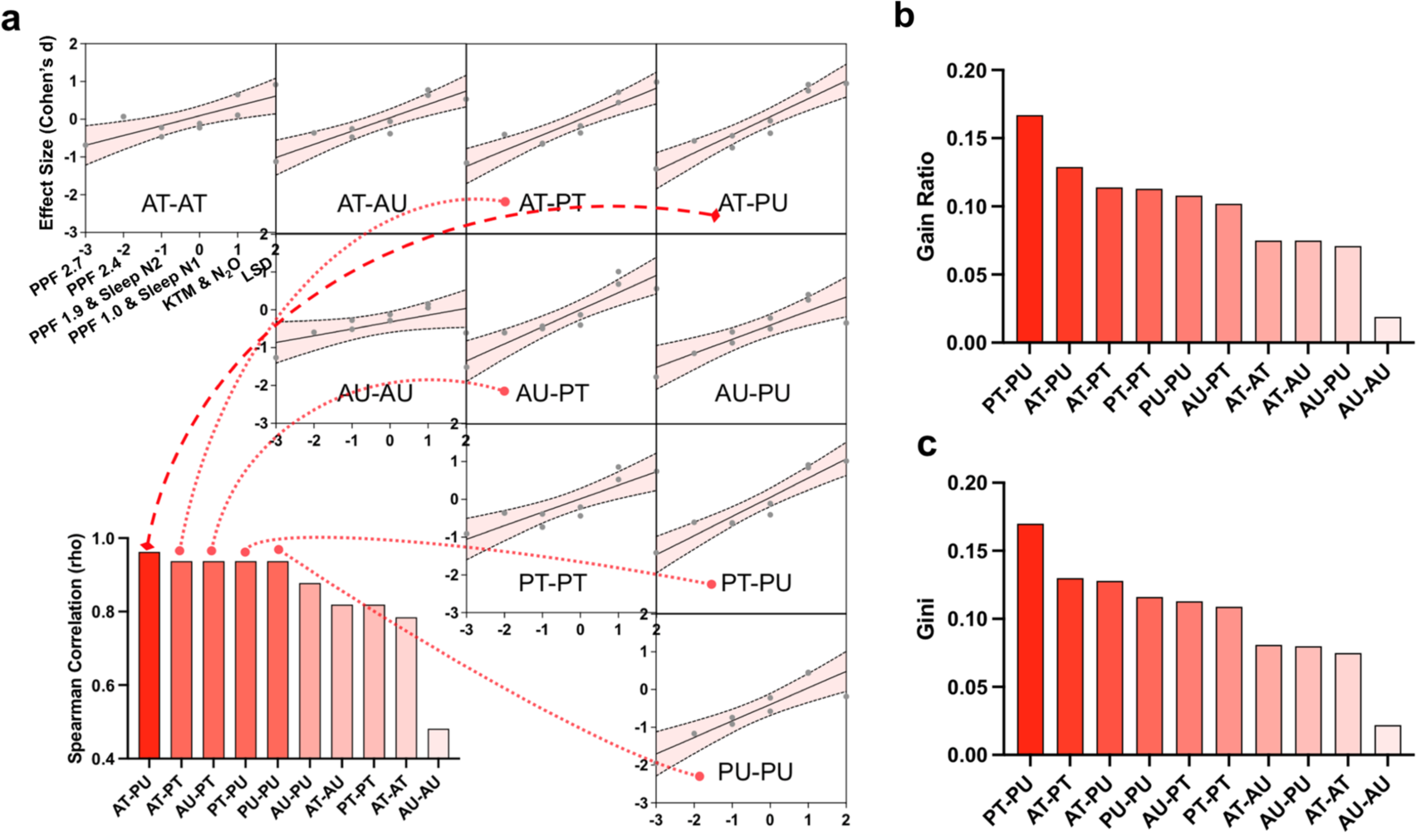
Correlation and Feature Importance of Brain Region Connectivity for Various States of Consciousness. **(a)** Spearman Correlation Analysis: Scatter plots show the rank correlation between different states of consciousness and functional connectivity effect sizes. Bar chart below highlights the Spearman correlation coefficients for each region pair. **(b)** Bar plot displays the feature ranking based on the gain ratio, a machine learning metric used to assess the importance of functional connectivity features in distinguishing enhanced consciousness (psychedelic states) from reduced consciousness (sleep and sedation). **(c)** Gini index, another feature importance metric.

Finally, to determine which type of functional connectivity (e.g., AT-PU or PT-PU) explains the greatest variance in the data across various states of consciousness, we conducted a non-parametric correlation analysis (Spearman correlation). We assigned -3 as deepest sedation (propofol 2.7 μg/ml), -2 as propofol 2.4 μg/ml, -1 as propofol 1.9 μg/ml and sleep N2, 0 as propofol 1.0 and sleep N1, 1 as ketamine and nitrous oxide, 2 as LSD (Jang et al., 2024a). Smaller positive values represent non-classical psychedelics, while the largest positive value represents classical psychedelic states. We correlated these state signs and magnitudes with the functional connectivity effect sizes across all states of consciousness. Our findings reveal that AT-PU explained the greatest variance of states of consciousness (rho = 0.96, pFDR = 0.001), followed by AT-PT, AU-PT, PT-PU, and PU-PU (rho = 0.94, pFDR = 0.001 for each). These results suggest that both A-P and P-P connections play a role in differentiating states of consciousness. In addition to correlation analysis, we employed feature ranking methods from machine learning to assess which features (i.e., functional connectivity between and within AT, AU, PT, and PU regions) best differentiate between enhanced consciousness states (psychedelic) and reduced consciousness states (sleep and sedation). We used two common feature importance metrics: gain ratio and Gini index. Both metrics identified PT-PU as the top-ranked feature, followed by AT-PU and AT-PT. This difference can be explained by the fact that Spearman correlation measures the strength of the monotonic relationship between functional connectivity and states of consciousness, highlighting AT-PU as the strongest in this regard. In contrast, feature ranking methods like gain ratio and Gini index assess the contribution of each connection to the overall classification task, where PT-PU demonstrated the highest importance in distinguishing between enhanced and reduced states of consciousness. However, the results are still consistent across these methods, as both highlight the critical role of A-P and P-P connections in differentiating states of consciousness, even though the specific rankings may vary slightly.

## Discussion

In this study, we systematically tested competing theories of consciousness by examining a broad spectrum of altered conscious states using a robust methodological framework. We demonstrated that psychedelic states of consciousness are linked to increased global functional connectivity and decreased local neural synchrony, while diminished states of consciousness during sleep and sedation displayed the opposite pattern. These findings suggest that consciousness arises from global brain network interactions rather than localized activity. This mirror-image pattern between enhanced and diminished states was observed in both anterior-posterior and within-posterior brain regions but not exclusively within the anterior part of the brain. Overall, the data (1) support global theories of consciousness in relation to varying states of consciousness, (2) do not support local theories of consciousness in either sensory or prefrontal cortex alone, (3) suggest that both anterior-posterior and posterior-posterior integration are crucial in defining various states of consciousness. Our findings also bridge the gap between GNW and IIT by highlighting the critical role of higher-order integrative cortical regions in consciousness.

Our study provides compelling empirical evidence for global theories of consciousness. While prior research has identified alterations in between-network and within-network integration during psychedelic states (Dai et al., 2023b; McCulloch et al., 2022; Shinn et al., 2023; Siegel et al., 2024), sleep (Horovitz et al., 2009; Tagliazucchi and Laufs, 2014), and sedation (Bonhomme et al., 2016; Boveroux et al., 2010; Huang et al., 2018; Jang et al., 2024b; Jordan et al., 2013; Palanca et al., 2015; Ranft et al., 2016), our work advances the field by simultaneously analyzing both global and local connectivity across a diverse spectrum of altered states. We demonstrate a striking mirror-image pattern: psychedelic states exhibit increased global functional connectivity and decreased local neural synchrony, while sleep and deep sedation show the opposite. This pattern, observed in both anterior-posterior and within-posterior brain regions, highlights the crucial role of global brain network interactions in supporting conscious experience. Our findings challenge the adequacy of local theories and bridge the gap between GNW and IIT, which may represent an important step forward in addressing the unresolved questions about the neural correlates of consciousness.

Moreover, we introduce a novel approach to the “front vs. back” debate of consciousness (Boly et al., 2017) by categorizing brain regions along a unimodal-transmodal axis, reflecting the functional hierarchy from basic sensory processing to higher-order cognitive integration. By examining connectivity within and between anterior transmodal, anterior unimodal, posterior transmodal, and posterior unimodal regions, we added a new degree of specificity in understanding the roles of lower-level sensory versus higher-order cognitive regions. We found that transmodal areas are more critical in A-P connectivity, while both transmodal and unimodal areas play key roles in P-P connectivity. This aligns with previous psychedelic research, which showed enhanced global functional connectivity in transmodal regions, such as the frontoparietal and default-mode networks during LSD experiences (Girn et al., 2022; Tagliazucchi et al., 2016). The focus on transmodal areas in GNW (Dehaene and Changeux, 2011; Dehaene and Naccache, 2001; Mashour et al., 2020) aligns with our data on connectivity between anterior and posterior transmodal regions. Within the framework of IIT (Tononi, 2008, 2004; Tononi et al., 2016), the inclusion of unimodal visual areas in the posterior “hot zone” helps explain the strong posterior connectivity observed when both transmodal and unimodal areas are considered. Although GNW underscores the importance of frontal regions, it emphasizes their dynamic interaction with posterior regions (Mashour et al., 2020). Our observation of a weaker effect of A-A connectivity suggests that frontal areas alone may be insufficient to sustain consciousness, further reinforcing this perspective.

We applied the measure of global efficiency (Jang et al., 2024b), calculated based on the inverse of shortest path length between regions, to assess the relative topological efficiency of information transfer between anterior and posterior brain regions, providing a deeper insight into network topology across states of consciousness. While the overall findings for functional connectivity and topological integration (i.e., global efficiency) are consistent, subtle differences emerge upon closer examination. For example, changes in efficiency were more strongly linked to within-posterior regions (i.e., PU-PU, PT-PU, and PT-PT). Topological integration, as opposed to functional connectivity, normalizes connectivity strength against a null model, highlighting the efficiency of information transfer independently of the raw connectivity values. Thus, functional connectivity and topological integration may provide complementary views on the brain’s functional organization.

Our rank correlation analysis revealed strong quantitative links between connectivity changes and levels of consciousness, with specific region pairs (e.g., AT-PU) showing the strongest correlations with altered states. Finally, feature importance analysis identified PT-PU as the top-ranked feature in distinguishing between enhanced (psychedelics) and reduced (sleep, sedation) consciousness. Together, these analyses provided a powerful quantitative framework for identifying the brain networks critical to altered consciousness. They revealed a mirror-image pattern, with heightened global integration during psychedelic states sharply contrasting with the diminished global integration observed during sleep and sedation, effectively capturing the opposing neural correlates of enhanced and reduced states of consciousness.

This study has limitations. First, it is important to acknowledge that increased connectivity, whether local or global, does not always equate to higher levels of consciousness. For example, excessive connectivity can lead to pathological states, such as seizures, where neural synchronization is heightened but consciousness is impaired. Second, the reliance on multiple datasets introduces heterogeneity, which may affect the generalizability of our results across different experimental settings or populations. Additionally, our results showed a decrease in ReHo at the highest doses of propofol, indicating reduced local connectivity. These doses of propofol approach anesthetic levels, going beyond mere unconsciousness and diverging from sleep-related patterns of increased local connectivity. Since deep sedation and sleep are more mechanistically similar than anesthesia and sleep, this divergence at higher propofol doses is expected (Franks and Wisden, 2021; Vanini et al., 2020). Anesthesia disrupts both global and local connectivity more profoundly, as supported by previous research (Huang et al., 2018). Lastly, although we analyzed both global and local connectivity, our results cannot fully support or invalidate local frameworks for consciousness, such as the Recurrent Processing Theory (Lamme et al., 2001; Lamme, 2010, 2006), which attempts to explain the mechanism for content of consciousness, particularly in studies that utilize visual tasks. Since our results are based on resting-state analysis, they cannot capture all the nuances of conscious processes.

Despite these limitations, our methods remain a valuable surrogate for coarse-grained tracking. By incorporating novel metrics like the topological integration and exploring the unimodal-transmodal axis, we offer new insights into the topological efficiency and hierarchical organization of brain networks during various states of consciousness. Our study advances the field by providing systematic support for global theories of consciousness and a clearer understanding of how large-scale brain networks and localized processes contribute to changes in consciousness. This work also bridges the gap between two competing theories, demonstrating that both anterior-posterior and within-posterior connectivity are critical for understanding various states of consciousness. Furthermore, our empirical findings suggest that different theories of consciousness may converge on the fundamental neuronal mechanisms and can be both compatible and complementary, despite marked differences in their theoretical foundations. By adopting a unifying, integration-oriented approach, our study combines valuable elements from various theories, providing a more comprehensive view of the neural underpinnings of consciousness.

## Methods

### Dataset 1: LSD

Fifteen healthy subjects (5 females; mean age ± standard deviation: 38.4 ± 8.6 years) were sourced from the OpenNEURO database (Carhart-Harris et al., 2016). The study design involved two experimental sessions, where participants received either a placebo or 75 μg of intravenous LSD in a counterbalanced order. Each participant completed three scans: the first and third were eyes-closed resting-state sessions, while the second involved a music listening task. Each scan duration was 7 minutes. This study utilized a 3T GE HDx scanner. For echo-planar imaging (EPI), the scan settings were as follows: 35 slices, a repetition time/echo of 2000/35 ms, 3.4 mm slice thickness, a 220 mm field of view, and a 90° flip angle. High-resolution T1 images were acquired as well.

In addition, only preprocessed data were available in the released dataset, followed by several steps. Initially, the first three volumes of each scan were removed to ensure stability. De-spiking was then conducted to correct signal artifacts, followed by slice time correction to align image acquisition timings. Motion correction was applied to counteract participant movement, and brain extraction isolated brain tissue from other elements in the images. The images were aligned to anatomical scans via rigid body registration and further aligned to a 2mm MNI brain template through non-linear registration. The dataset was then scrubbed using a FD threshold of 0.4, with a maximum of 7.1% of volumes scrubbed per scan, replacing them with the mean of surrounding volumes. Further processing included applying a 6mm kernel for spatial smoothing, filtering the data within the 0.01–0.1 Hz range, removing signal drifts through linear and quadratic de-trending, and eliminating motion- and anatomy-related artifacts through regression.

### Dataset 2: Ketamine

The research was approved by the Institutional Review Board of Huashan Hospital, Fudan University, and all participants provided written informed consent (Dai et al., 2023b; Huang et al., 2020). Twelve right-handed individuals (5 females; mean age ± standard deviation: 41.4 ± 8.6 years) were enrolled. Participants had no history of neurological disorders, significant organ dysfunction, or neuropsychiatric medication use, and were classified as American Society of Anesthesiologists physical status I or II. Intravenous ketamine was administered while fMRI scans were performed without interruption throughout the experiment, which spanned 44 to 62 minutes. A 10-minute baseline recording of the conscious state was conducted at the start, with the exception of two participants who had shorter baseline durations of 6 and 11 minutes. Ketamine was then administered at a rate of 0.05 mg/kg per minute over 10 minutes (total dose: 0.5 mg/kg), followed by an increased rate of 0.1 mg/kg per minute for another 10 minutes (cumulative dose: 1.0 mg/kg). Two participants only received the second dose. Once the ketamine infusion was complete, participants naturally regained consciousness. Our analysis focused on subanesthetic ketamine administration, which is associated with psychedelic experiences. This study utilized a 3T Siemens MAGNETOM scanner. For EPI, the scan settings were as follows: 33 slices, a repetition time/echo time of 2000/30 ms, 5mm slice thickness, a 210 mm field of view, a 64 × 64 image matrix, and a 90° flip angle. High-resolution T1 images were acquired as well.

### Dataset 3: Nitrous oxide

Sixteen healthy participants (8 females, mean age ± standard deviation: 24.6 ± 3.7 years) were recruited for this study, approved by the Institutional Review Board of the University of Michigan Medical School (HUM00096321), with all participants giving written informed consent (Dai et al., 2023b, 2023a). All participants were classified as American Society of Anesthesiologists physical status I, exclusion criteria included any history of drug abuse, psychosis, and other medical conditions as detailed in the trial registry (https://www.clinicaltrials.gov/ct2/show/NCT03435055). Two participants were excluded due to excessive head motion (affecting 50% of their fMRI data) and one due to incomplete scanning, leaving a final sample of 15 healthy subjects.

The experimental design consisted of two conditions: one conducted prior to and one during the administration of subanesthetic nitrous oxide (35%). Each condition included a 6-minute resting-state scan, a 3-minute visual task scan, and a 6-minute scan during a cuff-pain stimulus. This study utilized a 3T Philips Achieva scanner. For EPI, the scan settings were as follows: 48 slices, a repetition time/echo time of 2000/30ms, 3 mm slice thickness, a 200 mm field of view, and a 90° flip angle. High-resolution T1 images were acquired as well.

### Dataset 4: Sleep

Thirty-three healthy participants (16 females, mean age ± standard deviation: 22.1 ± 3.2 years) were recruited from the OpenNEURO database (Gu et al., 2023, 2022), with informed consent obtained from all participants. The dataset includes three non-REM sleep stages (N1, N2, and N3), in addition to an awake resting-state condition. These stages were identified using electroencephalogram signatures analyzed by a registered polysomnographic technologist. This study utilized a 3T Siemens Prisma scanner. For EPI, the scan settings were as follows: 35 slices, a repetition time/echo time of 2100/25 ms, 4 mm slice thickness, a 240 mm field of view, and a 90° flip angle. High-resolution T1 images were acquired as well. Only N1 (n=33) and N2 (n=29) were included in the analysis due to the limited number of subjects in N3 (n=3).

### Dataset 5: Propofol Sedation (1.0 and 1.9 μg/mL Doses)

Fifteen healthy participants (6 females, mean age ± standard deviation: 26.7 ± 4.8 years) were recruited for this study, approved by the Institutional Review Board of the Medical College of Wisconsin. All participants provided written informed consent, were classified as American Society of Anesthesiologists physical status I or II, and were scheduled for elective surgery to remove pituitary microadenomas. (Huang et al., 2018; Liu et al., 2017). Three participants were excluded due to excessive movement and MRI technical issues, leaving 12 participants for analysis.

Behavioral responsiveness was measured using the Observer’s Assessment of Alertness/Sedation (OAAS) scale. During baseline and recovery, participants were fully responsive to verbal cues, indicated by an OAAS score of 5. In the light sedation phase, participants responded lethargically to verbal commands, corresponding to an OAAS score of 4, while deep sedation was marked by the absence of a response, with OAAS scores ranging from 1 to 2. Individual propofol target plasma concentrations varied (light sedation: 0.98 ± 0.18 μg/mL; deep sedation: 1.88 ± 0.24 μg/mL), reflecting personal differences in sensitivity to the anesthetic. Propofol infusion rates were manually adjusted using STANPUMP to maintain steady sedation, balancing drug accumulation and elimination. Throughout the study, participants were monitored according to ASA standards, including electrocardiogram, blood pressure, pulse oximetry, and end-tidal CO2, with supplemental oxygen provided via nasal cannula. Resting-state data were collected across four 15-minute scans, each representing a different condition: baseline consciousness, light sedation, deep sedation, and recovery. This study utilized a 3T GE Signa 750 scanner. For EPI, the scan settings were as follows: 41 slices, a repetition time/echo time of 2000/25 ms, 3.5 mm slice thickness, a 224 mm field of view, and a 77° flip angle. High-resolution T1 images were acquired as well.

### Dataset 6: Propofol Sedation (2.4 μg/mL Doses)

Twenty-six healthy participants (13 females, mean age ± standard deviation: 25.0 ± 4.1 years) were recruited for this study, which was approved by the Institutional Review Board of the University of Michigan, and all participants provided written informed consent (Huang et al., 2023, 2021a, 2021b). All participants were classified as American Society of Anesthesiologists physical status I, exclusion criteria included any history of drug abuse, psychosis, and other medical conditions. One participant was excluded due to excessive movement, leaving 25 participants for analysis.

Before the study, participants fasted for eight hours. On the experiment day, a preoperative evaluation, including a physical exam, was conducted by an anesthesiologist. Two anesthesiologists continuously monitored vital signs, including breathing, heart rate, end-tidal CO2, pulse oximetry, and ECG. Noninvasive arterial pressure was recorded using an MR-compatible monitor. Participants received 0.5 mL of 1% lidocaine for local anesthesia before intravenous cannula insertion, and oxygen was delivered at 2 L/min through a nasal cannula. Propofol, chosen for its minimal impact on cerebral blood flow and precise titration capability, was administered via target-controlled bolus and infusion, based on the Marsh pharmacokinetic model using STANPUMP (http://opentci.org/code/stanpump). Dosages increased in 0.4 μg/mL increments until participants showed no behavioral response, with the target concentration maintained for an average of 21.6 ± 10.2 minutes before recovery.

Behavioral responsiveness was assessed by a rubber ball squeeze task, with responses quantified using the BIOPAC MP160 system and AcqKnowledge software. Sixty motor response trials, spaced 90 seconds apart, were conducted during scanning sessions. Between trials, participants engaged in mental imagery tasks such as imagining playing tennis or navigating a space. Further experimental details are available in prior publications.

This study utilized a 3T Philips scanner. For EPI, the scan settings were as follows: 28 slices, a repetition time/echo time of 800/25 ms (MB factor of 4), 4 mm slice thickness, a 220 mm field of view, and a 76°flip angle. High-resolution T1 images were acquired as well. Four fMRI sessions were conducted as part of the protocol: a 15-minute conscious baseline, a 30-minute session during and post-propofol infusion, followed by a 15-minute recovery baseline.

### Dataset 7: Propofol Sedation (2.7 μg/mL Doses)

Thirty healthy participants (20 females, mean age ± standard deviation: 24.4 ± 5.2 years) with complete scan data were included in this study, which was approved by the Institutional Review Board of the University of Michigan, and all participants provided written informed consent. All participants were classified as American Society of Anesthesiologists physical status I, exclusion criteria included any history of drug abuse, psychosis, and other medical conditions. Four participants were excluded due to excessive movement and MRI technical issues, leaving 26 participants for analysis.

The anesthetic procedure was similar to Dataset-6, with propofol manually adjusted to achieve effect-site concentrations of 1.5, 2.0, 2.5, and 3.0 μg/mL. Each level was held for 4 minutes to titrate the dosage and determine the threshold for loss of responsiveness (LOR). To minimize head motion artifacts, the concentration was maintained one step higher than the LOR threshold for about 32 minutes (e.g., if LOR occurred at 2.0 μg/mL, 2.5 μg/mL was maintained). In rare cases, if participants remained responsive at 3.0 μg/mL, the concentration was raised to a maximum of 4.0 μg/mL. The infusion was then stopped, and participants engaged in behavioral tasks, rest, or listened to music.

Eight fMRI scans, each lasting 16 minutes, were conducted over a 2.5-hour session. These included baseline scans (Rest1 and Music1), LOR scans (Rest2 and Music2), and recovery scans (Rest3 and Music3). Between each scan, participants had 1–5 minute breaks. Resting-state scans required participants to lie still with eyes closed, while music-listening involved tracks from Jazz, Rock, Pop, and Country genres, played in random order. During behavioral testing, participants were prompted to squeeze a rubber ball every 10 seconds for 96 cycles, following an audio cue delivered through headphones. Grip strength was measured using the BIOPAC MP160 system. Behavioral transitions during propofol administration were identified by missed and completed squeezes, marking the onset and recovery of responsiveness. This study utilized a 3T Philips scanner. For EPI, the scan settings were as follows: 40 slices, a repetition time/echo time of 1400/30 ms (MB factor of 4), 2.9 mm slice thickness, a 220 mm field of view, and a 76° flip angle. High-resolution T1 images were acquired as well.

### fMRI data preprocessing

For the preprocessing of fMRI data in this study, we utilized the AFNI software. The procedure encompassed several steps: First, the initial two frames of each scan were removed to ensure signal stability. This was followed by slice-timing correction to adjust for temporal differences in the acquisition of slices. Second, head motion correction and realignment were performed. Head motion was assessed using frame-wise displacement (FD), calculated as the Euclidean Norm of the six motion parameters. Frames where the FD exceeded 0.4mm, along with the preceding frame, were excluded from the analysis. Third, T1 anatomical images were coregistered for precise alignment, followed by spatial normalization into Talairach space (Talairach and Tournoux, 1988) and resampling to 3 mm isotropic voxels to standardize image coordinates. Fourth, time-censored data underwent band-pass filtering between 0.01–0.1Hz using AFNI’s 3dTproject. Simultaneously, linear regression was applied to eliminate unwanted components such as linear and nonlinear drift, head motion time series and its derivative, as well as mean time series from white matter and cerebrospinal fluid. Fifth, spatial smoothing (6mm Gaussian kernel) was performed. Finally, each voxel’s time series was normalized to zero mean and unit variance to ensure data consistency.

### Regional Homogeneity (ReHo) analysis

In our study, ReHo was computed to quantify the local synchronization of neural activity by examining the time-series similarity within clusters of neighboring voxels. The ReHo metric was derived using Kendall’s coefficient of concordance (Kendall’s W), which measures the degree of agreement among the ranks of time series data. The ReHo values for each voxel were calculated using the following formula:

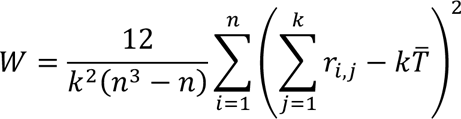

Here, *r_i, j_* signifies the rank of the ith time point in the jth voxel, *k* represents the number of voxels in the cluster, *n* is the number of time points, and 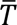. is the mean rank across all voxels and time points. The value of Kendall’s W ranges from 0, indicating no agreement, to 1, indicating complete agreement, thus reflecting the degree of local functional connectivity. The ReHo maps generated from this analysis provided insights into the patterns of functional connectivity under various conditions within our study, thereby enriching our comprehension of the brain’s functional organization. After the ReHo computation, we applied Fisher’s z-transformation to the ReHo values. This transformation standardizes the variance, facilitating more accurate correlation studies and comparisons across different subjects and conditions.

### Between-Network Functional Connectivity

To assess between-network functional connectivity, we utilized a well-established parcellation scheme (Yeo et al., 2011) to define 400 regions of interest (ROIs) across the brain, encompassing seven predefined functional networks: the default mode, frontoparietal, limbic, ventral attention, dorsal attention, somatomotor, and visual networks. Time series data were extracted from each ROI and used to compute Pearson correlation coefficients, generating a 400 x 400 functional connectivity matrix. For each of the seven networks, between-network connectivity was quantified by calculating the Pearson correlation between the ROIs within the given network and those in each of the other six networks. The resulting six between-network correlation values were averaged to produce a single between-network connectivity value for each network. Finally, the global between-network FC for the entire brain was obtained by averaging these network-specific values, providing an overall index of inter-network connectivity across the brain. Fisher’s z-transformation was applied to the correlation coefficients to normalize the data and ensure suitability for statistical analyses.

### Anterior and Posterior Functional Connectivity

To analyze anterior and posterior functional connectivity, we segmented the brain into two regions using the central sulcus as the anatomical boundary. The anterior region, which includes the frontal cortex, consisted of 180 out of the 400 ROIs, while the posterior region, encompassing the parietal, temporal, and occipital cortices, contained the remaining 220 ROIs. Time series data were extracted from each ROI, and functional connectivity was assessed across the regions within and between the anterior and posterior areas. Specifically, these calculations were performed for connectivity within the anterior region (A-A), between the anterior and posterior regions (A-P), and within the posterior region (P-P). Pearson correlation was applied to compute the relationships between ROIs, followed by Fisher’s z-transformation for statistical normalization.

### Anterior/Posterior vs. Transmodal/Unimodal functional connectivity

Using a gradient-based approach, we assessed functional connectivity across transmodal (higher-order cognitive) and unimodal (sensory-motor) areas within the anterior and posterior regions of the brain. This method relies on variations in functional connectivity strengths between brain regions. Specifically, it calculates a functional connectivity matrix that captures correlations between the time series data of different brain areas. Gradients are then extracted using dimensionality reduction techniques, such as diffusion mapping embedding, which identify gradual shifts in connectivity patterns across the brain. These gradients organize brain regions along a continuum from lower-level sensory-motor (unimodal) areas to higher-order cognitive (transmodal) regions. The gradient values were calculated from data of healthy adult subjects obtained through the Human Connectome Project by the BrainSpace toolbox. (https://brainspace.readthedocs.io/en/latest/). ROIs were categorized based on their gradient values, where ROIs with a gradient value greater than 0 were classified as transmodal, and those with a value less than 0 were classified as unimodal. To perform this analysis, we divided the ROIs into four groups: (1) anterior transmodal (AT) regions, comprising ROIs located in the anterior region (frontal cortex) with gradient values greater than 0; (2) anterior unimodal (AU) regions, comprising ROIs in the anterior region with gradient values less than 0; (3) posterior transmodal (PT) regions, consisting of ROIs in the posterior region (parietal, temporal, and visual cortices) with gradient values greater than 0; and (4) posterior unimodal (PU) regions, containing ROIs in the posterior region with gradient values less than 0. Time series data were extracted from these ROIs, and correlation matrices were computed to evaluate functional connectivity within and between the AT, AU, PT, and PU regions. This approach allowed for the analysis of functional connectivity patterns both along the anterior-posterior axis and across transmodal-unimodal regions.

### Functional Connectivity Density Analysis

To evaluate the density of functional connectivity between anterior-posterior and transmodal-unimodal brain regions, we computed the proportion of strong functional connections (defined as |effect size| > 0.8) for each pair of regions. Connectivity density was calculated by dividing the number of robust functional connections by the total number of possible connections within and between anterior transmodal (AT), anterior unimodal (AU), posterior transmodal (PT), and posterior unimodal (PU) regions. This allowed us to quantify changes in connectivity density across different consciousness states, including psychedelic, sleep, and sedative conditions. The resulting connectivity density values were visualized using circular plots, where lines between regions represented strong connections with effect sizes exceeding 0.8, providing a clear depiction of altered connectivity patterns under different conditions.

### Cognitive Term Word Cloud Analysis

To explore the cognitive functions associated with the identified brain regions, we used a meta-analytic approach leveraging data from Neurosynth (https://www.neurosynth.org/). For each region (AT, AU, PT, PU), we generated meta-analytic maps based on functional activation patterns, and correlated these maps with the corresponding anatomical masks of each region. The resulting correlation coefficients (r values) were used to identify cognitive terms most strongly associated with each region. Word clouds were generated, with the font size of each cognitive term proportional to its correlation coefficient. This visual representation highlighted the cognitive functions most relevant to each brain region in the context of the state of consciousness. A triangle beneath the word clouds was used to indicate the relationship between font size and the correlation strength, providing an intuitive understanding of the cognitive roles of different brain regions.

### Normalized multi-level global efficiency

We calculated the normalized multi-level global efficiency to assess the topological integration of the network independently of mere connectivity strength. The global efficiency of a binary network is defined as the average of the inverse shortest path lengths between all pairs of regions within the network:

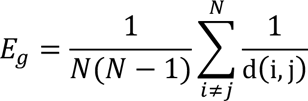

where N is the number of regions (nodes) in the network, and d(i,j) is the shortest path length between regions i and j. The shortest path length d(i,j) represents the minimal number of connections required to transfer information from region i to region j.

To address the weighted nature of the functional connectivity matrices, we adopted a multi-level thresholding methodology (Jang et al., 2024b). The Fisher z-transformed functional connectivity matrix was binarized at multiple thresholds t ranging from 0 to *r*_max_ in increments of 0.01, where *r*_max_ is the largest element in the functional connectivity matrix. For each thresholded binary matrix, we calculated the global efficiency. The multi-level efficiency *E*_*ML*_ was then determined by calculating the area under the curve of the global efficiency as a function of the threshold:

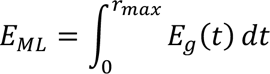

We normalized the multi-level efficiency by dividing it by the multi-level efficiency of a corresponding random null model 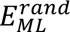:

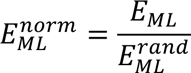

The random null model was generated by randomly rewiring the edges while preserving the degree distribution, using a custom-modified version of the ‘randmio_und’ function from the Brain Connectivity Toolbox (https://doi.org/10.1016/j.neuroimage.2009.10.003).

Between-network efficiency was calculated in a similar manner. When selecting pairs of regions for efficiency calculations, we only considered between-network pairs, of which the two regions belonged to different networks. Averaging was only performed over these between-network pairs. Additionally, when generating the random null model for between-network efficiency, we shuffled only the between-network edges, preserving the within-network connectivity structure.

### Rank Correlation Analysis of Functional Connectivity Across Conscious States

To investigate the relationship between functional connectivity and changes in conscious states, we performed a group-level non-parametric rank correlation (Spearman correlation) analysis between the effect size of changes in functional connectivity and the level of states of consciousness. States of consciousness were ranked as follows: the most negative value (-3) represented deep sedation under propofol 2.7 μg/mL, followed by -2 for propofol 2.4 μg/mL, and -1 for propofol 1.9 μg/mL and sleep N2. A value of 0 was assigned to Sleep N1 and propofol 1.0 μg/mL. Positive values were assigned to psychedelic states, with a rank of 1 corresponding to non-classical psychedelics (e.g., ketamine and nitrous oxide), and a rank of 2 corresponding to classical psychedelics (e.g., LSD). Spearman’s correlation coefficient (rho) was computed.

### Feature Ranking of Functional Connectivity for States of Consciousness Classification

To identify the most important functional connectivity features that differentiate between enhanced states of consciousness (e.g., psychedelic) and reduced states (e.g., sleep and sedation), we applied feature ranking techniques from machine learning. Two widely used feature importance metrics, gain ratio and Gini index, were employed to rank the z-scores of changes in functional connectivity. Functional connectivity features were derived from the correlation matrices computed for defined ROI pairs. These features, representing the strength of connectivity between specific brain regions, were used as input for the ranking process. Gain ratio, a normalized version of information gain, measures the relevance of each feature by evaluating how much information about the target variable (state of consciousness) is gained by splitting on that feature. The Gini index, commonly used in decision trees, evaluates feature importance based on how well it separates the different states of consciousness. By applying these feature ranking methods, we identified which functional connectivity features, within and between the anterior transmodal (AT), anterior unimodal (AU), posterior transmodal (PT), and posterior unimodal (PU) regions, contributed the most to distinguishing between different levels of consciousness.

### Statistics and Reproducibility

For all functional connectivity (FC) and regional homogeneity (ReHo) analyses, paired t-tests were conducted to compare each condition (psychedelic, sleep, and sedation) to its respective baseline. Statistical significance was determined using FDR-corrected p-values (p < 0.05) to account for multiple comparisons. Effect sizes were calculated using Cohen’s d for paired samples. All statistical tests used in this study were two-sided.

Spearman’s correlation was applied to assess the relationship between ranked states of consciousness and functional connectivity changes. FDR correction was used to ensure significance at p < 0.05 across all comparisons.

## Data availability

All data supporting the findings of this study are provided in Supplementary Data. The natural sleep fMRI dataset is available from OpenNEURO (https://openneuro.org/datasets/ds003768/versions/1.0.11). The LSD dataset is published at Openneuro (doi: 10.18112/openneuro.ds003059.v1.0.0). Access to additional data are not openly available due to reasons of participant privacy and are available from the corresponding author upon reasonable request. Data are located in controlled access data storage at University of Michigan Medical School.

## Code availability

Publicly available software and toolbox used for analyses include AFNI (https://afni.nimh.nih.gov/), JASP (v0.16.3; https://jasp-stats.org/), BrainSpace toolbox (https://brainspace.readthedocs.io/en/latest/), Brain Connectivity Toolbox (https://doi.org/10.1016/j.neuroimage.2009.10.003) and MATLAB R2022a (https://www.mathworks.com/products/new_products/release2022a.html)

## Acknowledgements

This work was funded by National Institutes of Health (Bethesda, Maryland, USA) grants R01-GM103894 (to A.G.H. and Z.H.), R01-GM111293 (to G.A.M., R.E.H.) and T32-GM103730 (to G.A.M., PI, and R.D., Z.H., Fellows).

## Authorship contribution statement

Conceptualization: R.D., Z.H., G.A.M. Methodology: R.D., Z.H., G.A.M. Investigation: R.D., Z.H., G.A.M. Data analysis and visualization: R.D., Co-data analysis Z.H. and H.J., Supervision: G.A.M, A.G.H., Writing—original draft: R.D., Writing—review & editing: All authors.

## Supplementary Information

**Figure S1.**
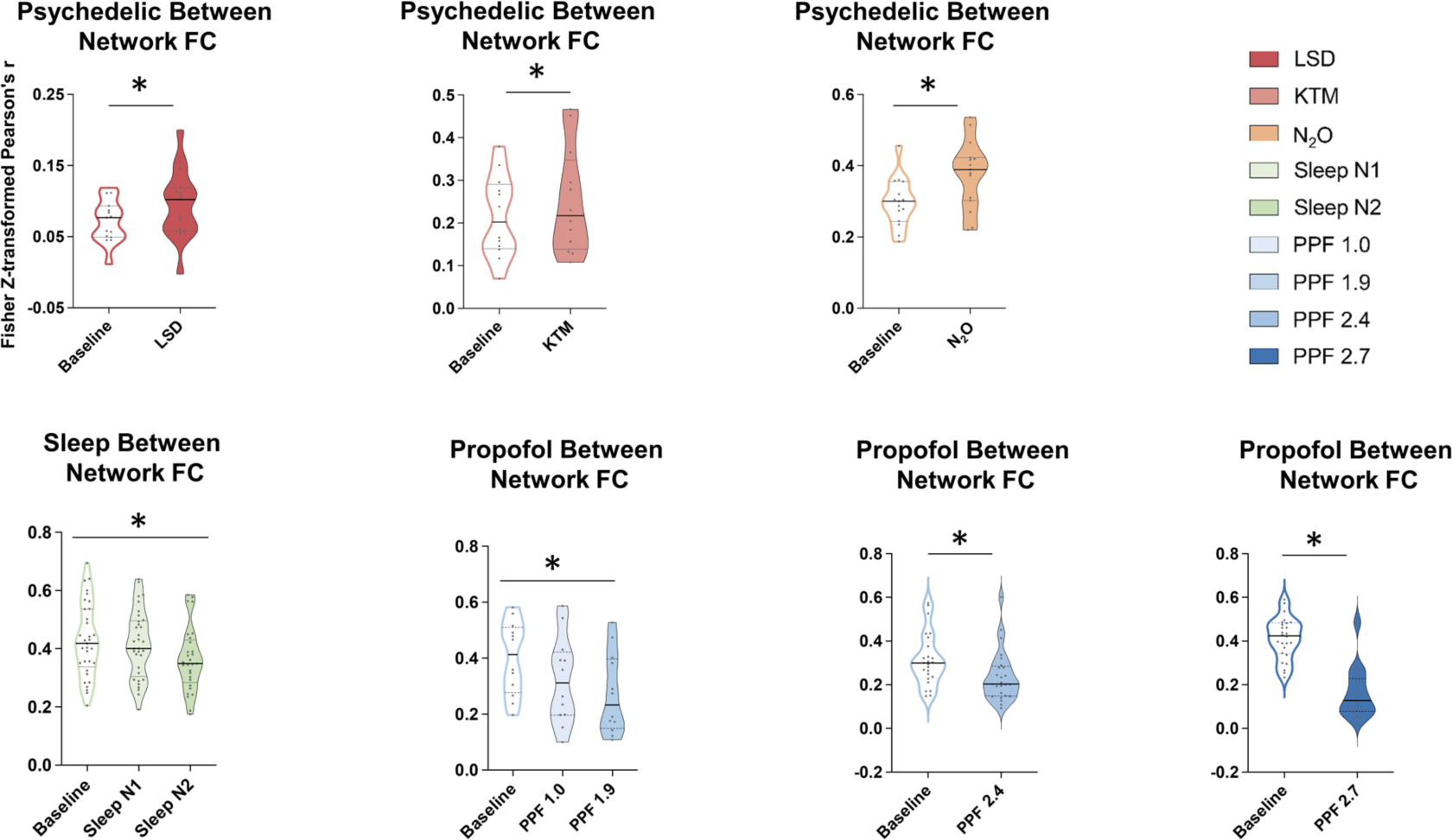
Between-Network Functional Connectivity Across States of Consciousness. Violin plots show between-network functional connectivity (FC; Fisher z-transformed Pearson’s r) for psychedelic states (LSD, KTM, N_2_O), sleep stages (N1, N2), and propofol-induced sedation at varying effect-site concentrations. Significant differences from baseline are marked by asterisks (*, FDR-corrected p < 0.05). LSD: lysergic acid diethylamide, KTM: ketamine, N_2_O: nitrous oxide, Sleep N1/N2: non-REM sleep stage 1/ stage 2, PPF 1.0: propofol 1.0 μg/ml, PPF 1.9: propofol 1.9 μg/ml, PPF 2.4: propofol 2.4 μg/ml, PPF 2.7: propofol 2.7 μg/ml.

**Figure S2.**
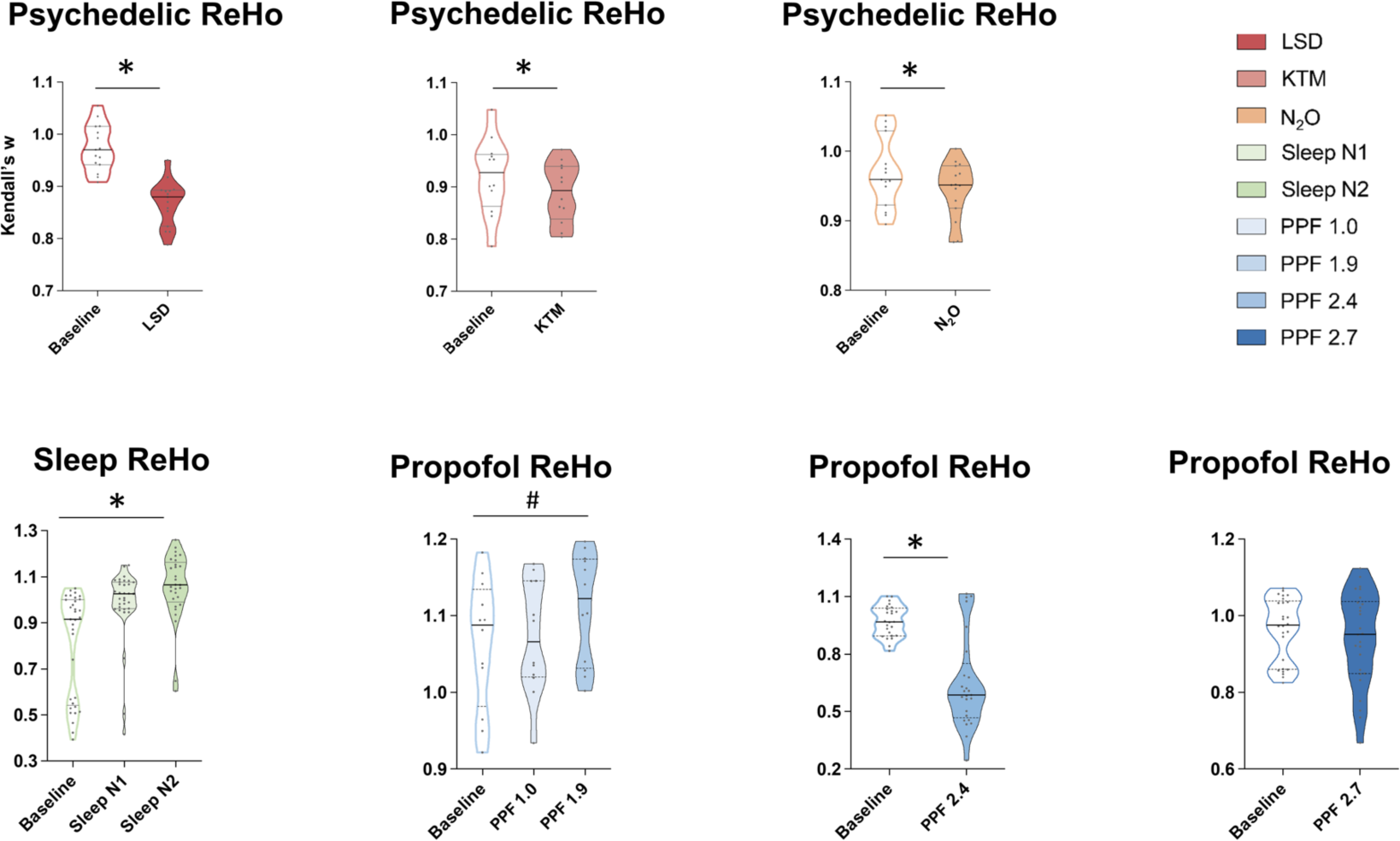
Regional Homogeneity (ReHo) Across States of Consciousness. Violin plots depict ReHo values (Kendall’s W) for psychedelic states (LSD, KTM, N_2_O), sleep stages (N1, N2), and propofol-induced sedation at varying effect-site concentrations. Significant differences from baseline are marked by asterisks (*, FDR-corrected p < 0.05), and significant differences before FDR correction are indicated by a hash (#, uncorrected p < 0.05). LSD: lysergic acid diethylamide, KTM: ketamine, N_2_O: nitrous oxide, Sleep N1/N2: non-REM sleep stage 1/ stage 2, PPF 1.0: propofol 1.0 μg/ml, PPF 1.9: propofol 1.9 μg/ml, PPF 2.4: propofol 2.4 μg/ml, PPF 2.7: propofol 2.7 μg/ml.

**Figure S3.**
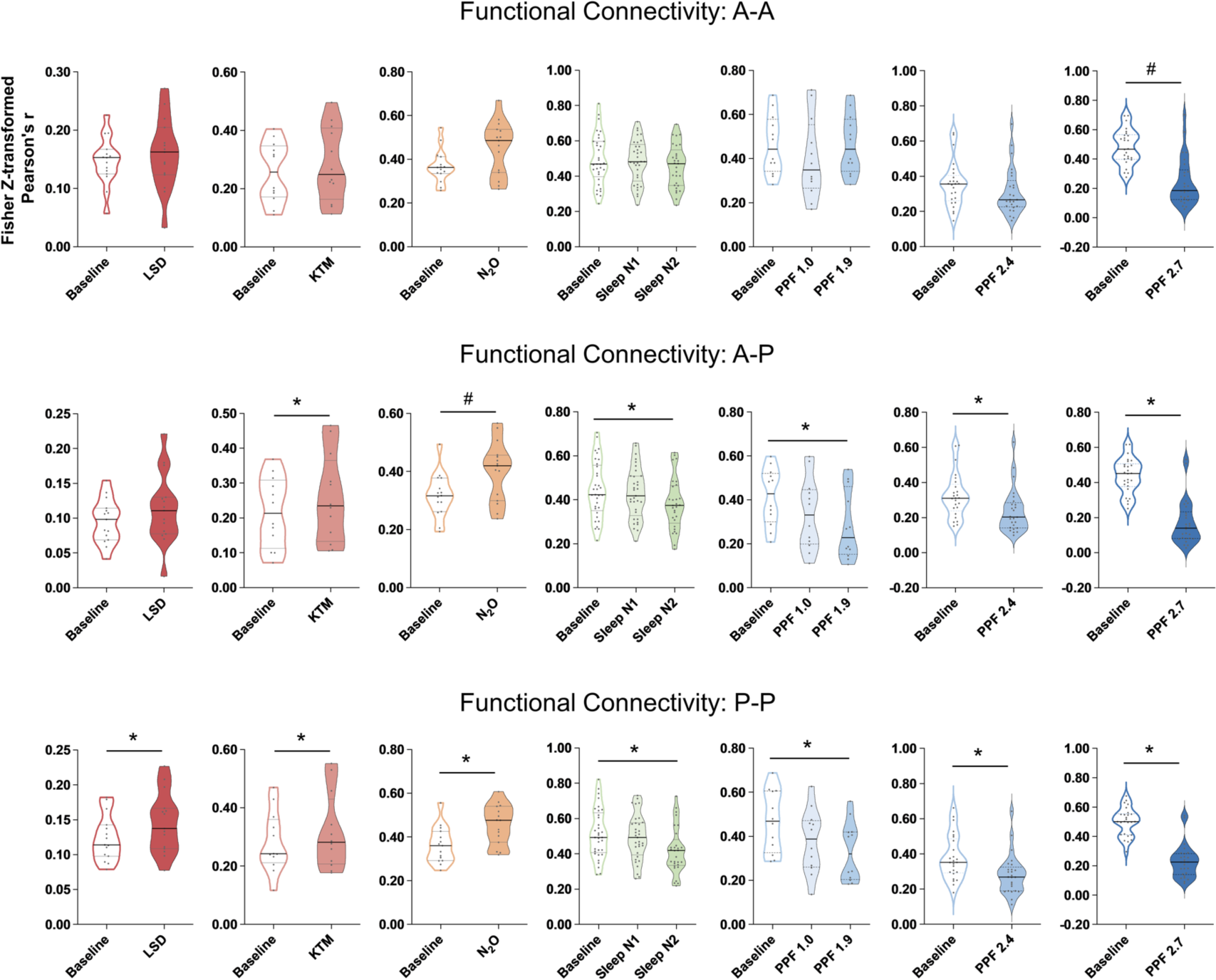
Functional Connectivity Within and Between Anterior and Posterior Regions in Different States of Consciousness. Functional connectivity was assessed within anterior regions (A-A), within posterior regions (P-P), and between anterior and posterior regions (A-P). Violin plots show functional connectivity (Fisher z-transformed Pearson’s r) for psychedelic states (LSD, KTM, N_2_O), sleep stages (N1, N2), and propofol-induced sedation at varying effect-site concentrations. Significant differences from baseline are marked by asterisks (*, FDR-corrected p < 0.05; #, uncorrected p < 0.05). LSD: lysergic acid diethylamide, KTM: ketamine, N_2_O: nitrous oxide, Sleep N1/N2: non-REM sleep stage 1/ stage 2, PPF 1.0: propofol 1.0 μg/ml, PPF 1.9: propofol 1.9 μg/ml, PPF 2.4: propofol 2.4 μg/ml, PPF 2.7: propofol 2.7 μg/ml.

**Figure S4.**
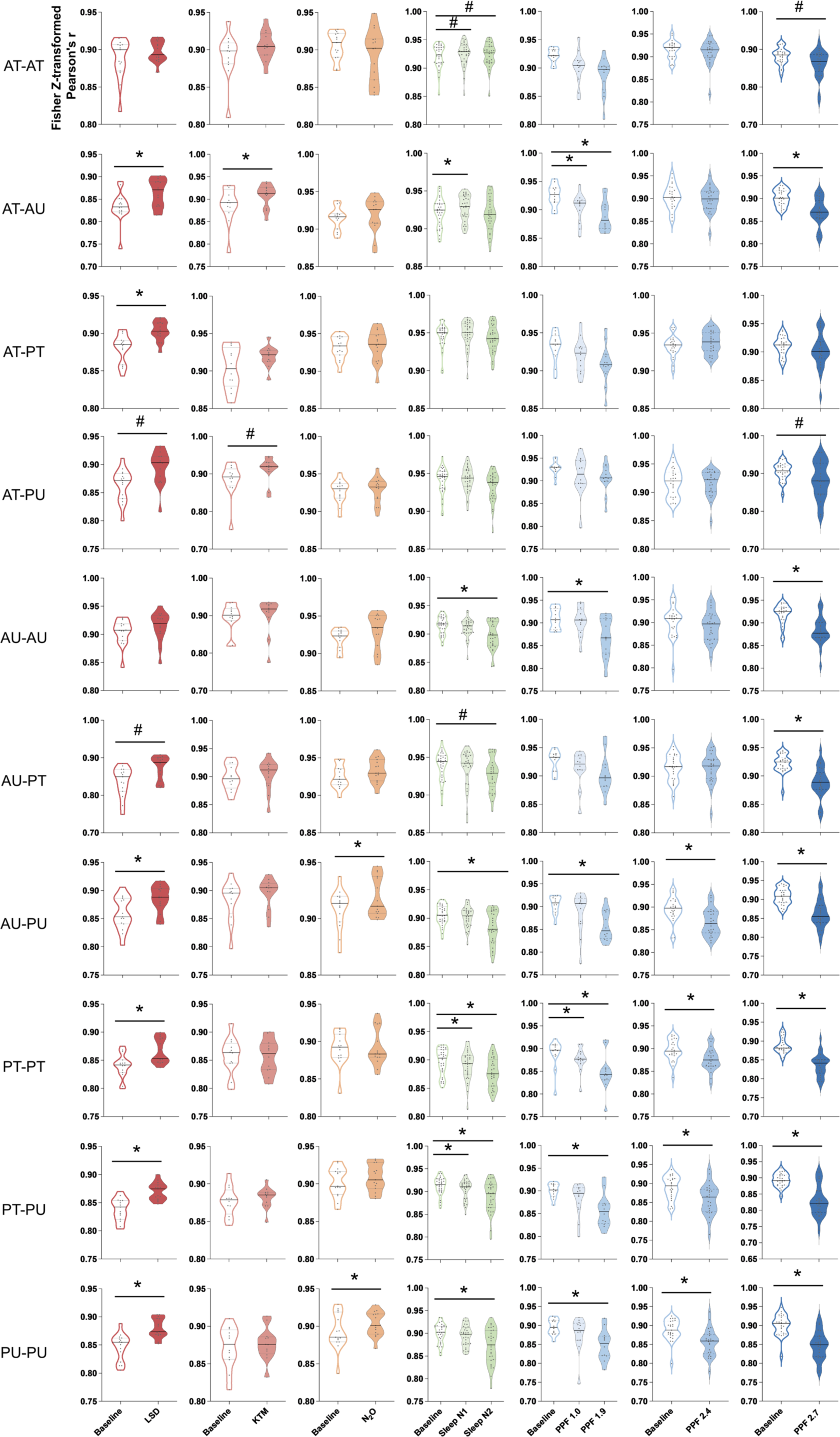
Functional Connectivity Within and Between Anterior/Posterior and Transmodal/Unimodal Regions in Different States of Consciousness. Functional connectivity was assessed between and within anterior transmodal (AT), anterior unimodal (AU), posterior transmodal (PT), and posterior unimodal (PU) regions. Violin plots show functional connectivity (Fisher z-transformed Pearson’s r) for psychedelic states (LSD, KTM, N_2_O), sleep stages (N1, N2), and propofol-induced sedation at varying effect-site concentrations. Significant differences from baseline are marked by asterisks (*, FDR-corrected p < 0.05; #, uncorrected p < 0.05). LSD: lysergic acid diethylamide, KTM: ketamine, N_2_O: nitrous oxide, Sleep N1/N2: non-REM sleep stage 1/ stage 2, PPF 1.0: propofol 1.0 μg/ml, PPF 1.9: propofol 1.9 μg/ml, PPF 2.4: propofol 2.4 μg/ml, PPF 2.7: propofol 2.7 μg/ml.

**Figure S5.**
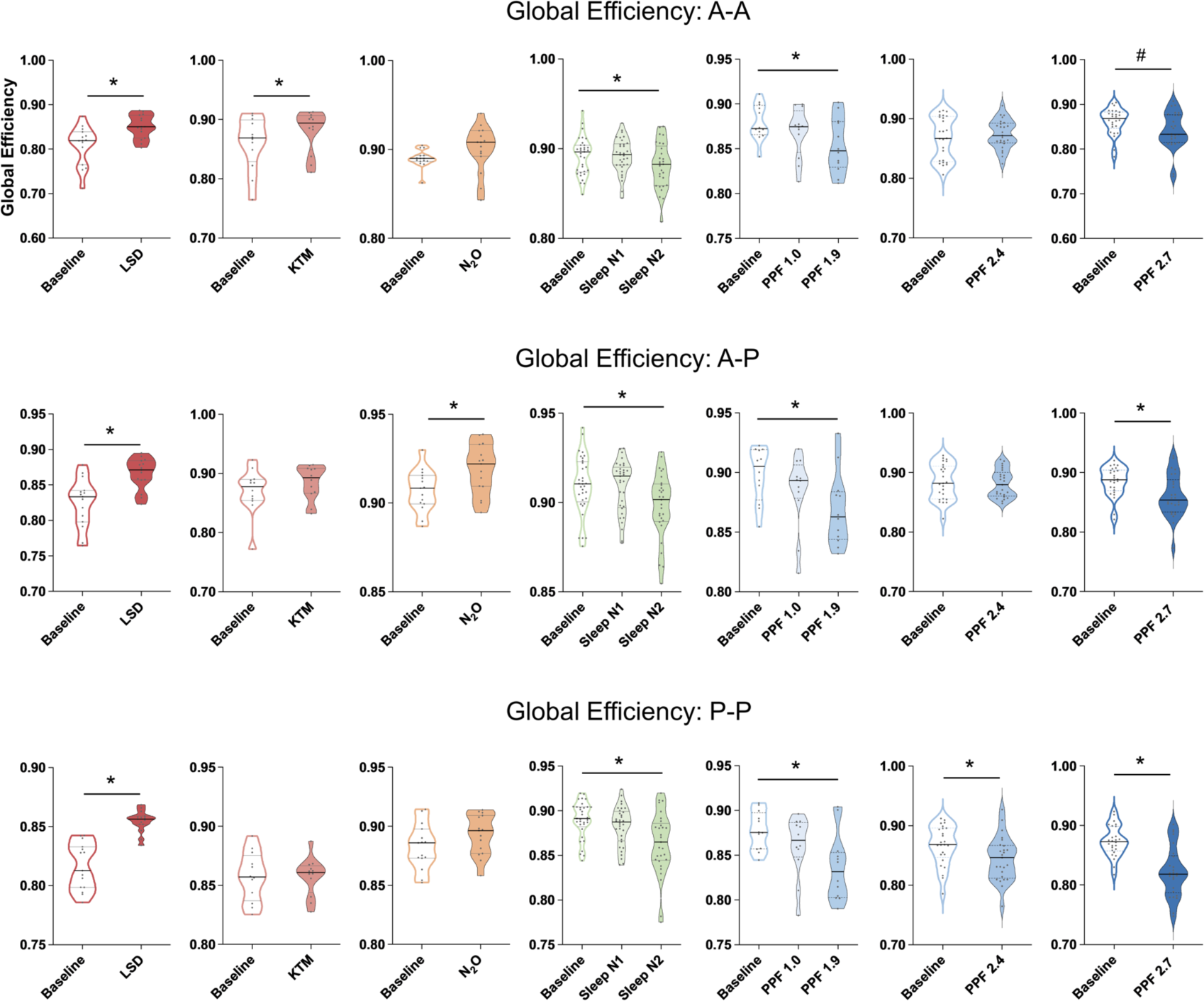
Global Efficiency Within and Between Anterior and Posterior Regions in Different States of Consciousness. Global efficiency was assessed within anterior regions (A-A), within posterior regions (P-P), and between anterior and posterior regions (A-P). Violin plots show functional connectivity (Fisher z-transformed Pearson’s r) for psychedelic states (LSD, KTM, N_2_O), sleep stages (N1, N2), and propofol-induced sedation at varying effect-site concentrations. Significant differences from baseline are marked by asterisks (*, FDR-corrected p < 0.05; #, uncorrected p < 0.05). LSD: lysergic acid diethylamide, KTM: ketamine, N_2_O: nitrous oxide, Sleep N1/N2: non-REM sleep stage 1/ stage 2, PPF 1.0: propofol 1.0 μg/ml, PPF 1.9: propofol 1.9 μg/ml, PPF 2.4: propofol 2.4 μg/ml, PPF 2.7: propofol 2.7 μg/ml.

**Figure S6.**
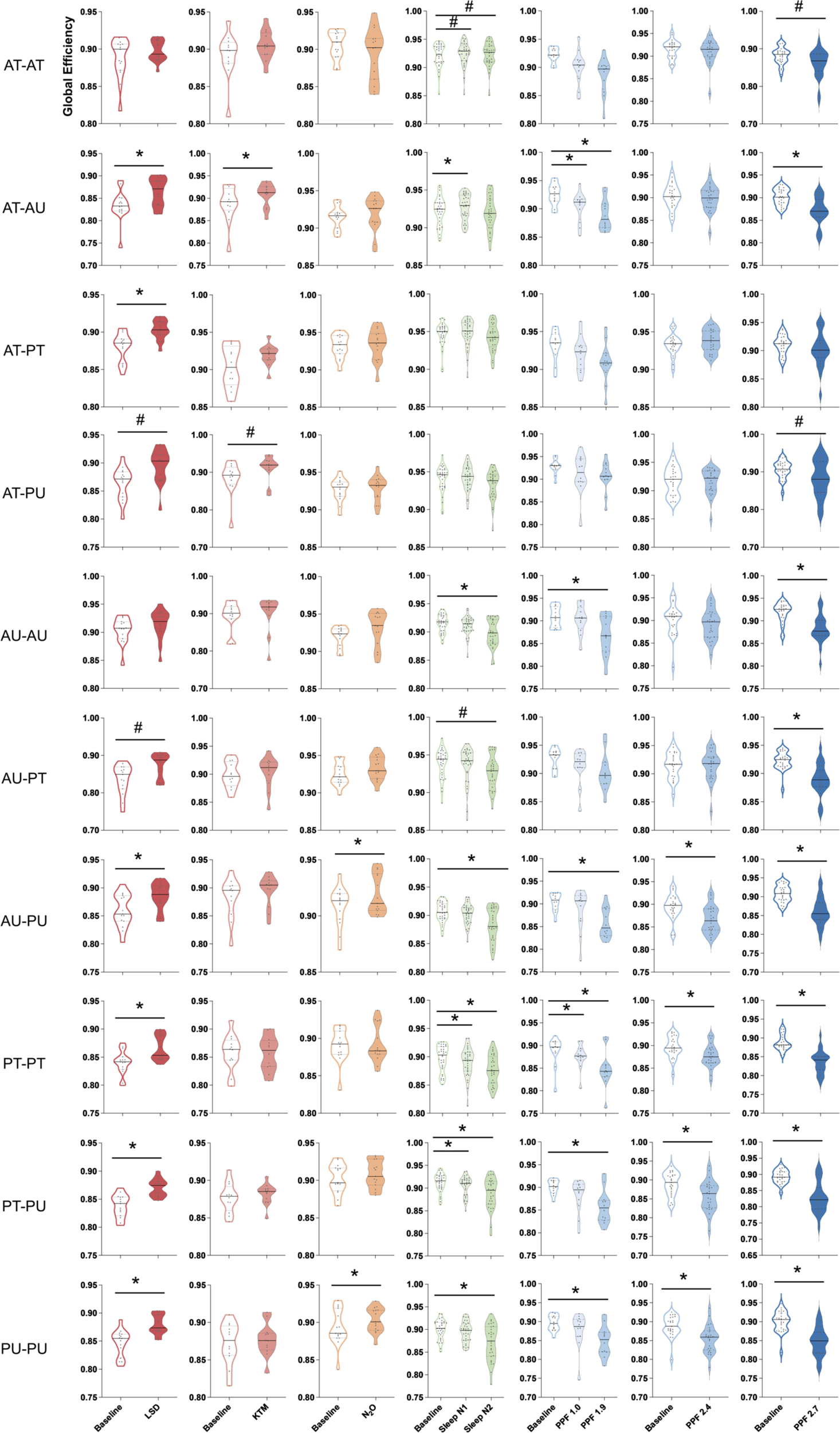
Global Efficiency Within and Between Anterior/Posterior and Transmodal/Unimodal Regions Across States of Consciousness. Global efficiency was assessed between and within anterior transmodal (AT), anterior unimodal (AU), posterior transmodal (PT), and posterior unimodal (PU) regions. Violin plots show functional connectivity (Fisher z-transformed Pearson’s r) for psychedelic states (LSD, KTM, N_2_O), sleep stages (N1, N2), and propofol-induced sedation at varying effect-site concentrations. Significant differences from baseline are marked by asterisks (*, FDR-corrected p < 0.05; #, uncorrected p < 0.05). LSD: lysergic acid diethylamide, KTM: ketamine, N_2_O: nitrous oxide, Sleep N1/N2: non-REM sleep stage 1/ stage 2, PPF 1.0: propofol 1.0 μg/ml, PPF 1.9: propofol 1.9 μg/ml, PPF 2.4: propofol 2.4 μg/ml, PPF 2.7: propofol 2.7 μg/ml.

